# Blank Spectrum Correction as a Robust Solution to Artifacts in Quantitative X-ray Fluorescence Mapping

**DOI:** 10.64898/2026.01.18.700193

**Authors:** Andrew M. Crawford, Julia Balough, Yu-Ying Chen, Qiaoling Jin, Keith W. MacRenaris, Seth Garwin, Teresa K. Woodruff, Chris Jacobsen, James E. Penner-Hahn, Thomas V. O’Halloran

## Abstract

X-ray fluorescence microscopy (XFM) continues to develop as a powerful quantitative technique for high resolution, label-free, elemental mapping of biological, environmental, and material samples. Methods for rigorously fitting spectra, increasing throughput, accounting for background signals, and deconvoluting overlapping emission lines continue to evolve. We show here that quantitative fits of XFM data obtained after removing a baseline, calculated by connecting peak edges, can be unexpectedly dependent upon acquisition dwell-time and spectral aggregation leading to differences in apparent elemental content. Using mouse preimplantation embryos and ovarian follicles as model samples, we demonstrate how these variables influence quantitative comparisons between samples. We find that subtracting an empirically measured blank spectrum instead of a baseline provides quantitative XFM elemental mapping results that are independent of dwell time and spectral aggregation dependencies.

---

Synchrotron-based X-ray fluorescence (XRF) microscopy (XFM) is a powerful technique for quantitatively mapping elemental distributions within cells and tissues on length scales ranging from nanometers to millimeters.^1–10^ With improving sources of synchrotron radiation, readout electronics, and detectors,^11–12^ XFM is shedding new light on diverse biological topics including, metal homeostasis, mammalian^13–20^ and amphibian^21^ egg fertilization and maturation, neurodegenerative disorders,^9, 22–24^ drug development,^24^ and bionanotechnology,^3^ among many others. The major advantages of XFM over other elemental imaging techniques are that it can be non-destructive (depending on source intensity and focusing optics), requires less chemical perturbation during sample preparation, has good detection limits for a variety of elements, and is subject to minimal matrix effects

XFM instruments typically focus intense X-ray beams with spot sizes from microns to nanometers^25^ on to a sample which is raster scanned through the beam path at user designated x- and y-coordinates and dwell times. XRF occurs when a photon, absorbed by an atom, causes ejection of a core shell electron followed by electronic relaxation and subsequent emission of an X-ray photon with an energy that correlates with atomic number. The full emission spectrum at each pixel is collected using an energy dispersive X-ray detector and subsequently fitted using a variety of programs^26–32^ to determine the element-specific signal at each pixel. By comparison to measurements from reference standards, this fitted data is then converted to quantitative two-dimensional elemental maps of the sample. One challenge associated with quantitative mapping of biological samples is the substantial peak overlap for samples containing dozens of elements present in quantities that span orders of magnitude. Most X-ray emissions have a natural line width of a few eV; however, the best energy dispersive X-ray detectors are resolution limited to ca. 150 eV.^33^ As a result, spectra must be fitted to deconvolute the peak overlap^34^ and account for a variety of contributions to the background. For XRF, the detected signal (F), either from an individual pixel or aggregate spectrum, contains fluorescence from the sample (S), a continuum (C) of diffuse scatter, and contributions from background fluorescence (B), such that F = S + C + B.^35^ The most widely used approach to isolate signal S is to first remove C using baseline subtraction.^36–41^ Baseline subtraction of XRF accounts for C by calculating a smooth baseline (L) which connects the leading and trailing edges of each peak, such that the residual F_SB_ = S + B can be calculated and then fit. After the data are fit, B is accounted for by selecting an off-sample area, calculating its average for each fitted element, and then subtracting that average value from the corresponding fitted element’s map (or aggregate ROI value). Alternatively, blank spectrum correction^27, 42^ accounts for both C and B by taking the mean fluorescence spectrum of the same off sample area and subtracting that spectrum instead of the baseline from the spectrum to be fitted, such that the residual F_S_ = S can be calculated and then fit.

In the process of comparing the outputs from blank spectrum correction^27, 42^ and baseline subtraction^32^, we have identified systematic concentration variations from baseline subtraction that are dependent on the dwell time and spectral aggregation. At the short exposure times necessary for mapping, per-pixel estimates of C have such low photon count values that baselines inaccurately model C, and due to peak overlap no amount of signal aggregation (i.e., adding pixels together) can produce a signal from which an accurate model of C can be derived.

This work is motivated by XFM studies of dynamic changes in essential element concentrations in mouse oocytes undergoing maturation in ovarian follicles^43^, and later as they undergo egg-to-embryo transitions after fertilization.^44^ In the course of these studies, we found that fitted values for baseline-corrected data sets unexpectedly depended on both dwell time and signal aggregation performed prior to fitting. We collected XFM data at multiple dwell times and compared various methods for fitting the data. We find that quantitation changes when either the dwell time of data collection changes or when fitting is done on the aggregate spectra of image regions of interest (ROI) as opposed to fitting individual pixel spectra and summing the results. Using both acquired spectra and simple simulations, we establish that these effects are a consequence of using baseline subtraction prior to the fitting of spectra and show that correction by an empirically measured blank spectrum can overcome this problem. Depending upon the element, we find the difference in quantitation can be small or quite large compared to the biological variation from sample to sample.

## Results

Comparison of aggregate and per-pixel spectra gives significantly different results for baseline subtraction but identical results for blank spectrum correction. XFM data for late blastocyst-stage mouse embryos were collected at beamline 2-ID-E of the Advanced Photon Source (APS, Argonne National Laboratory, Lemont, Illinois, USA). Whole blastocyst regions of interest (ROIs) were assigned manually for each scan using the fitted phosphorus map. Quantitative analyses using both blank spectrum correction and baseline subtraction were performed on the individual pixel spectra and on the aggregate spectrum of each ROI (**Fig. 1**, **Table 1**, **Table S1**, **Table S2**). Quantitative fitting from blank spectrum correction of per-pixel and aggregate spectra yielded identical answers within computational precision (**Table S2**); however, when baseline subtraction was used, aggregate spectrum fitting always resulted in lower concentrations and narrower distributions than per-pixel fitting (**Table 1a**), and neither agreed with their blank spectrum correction counterpart (discussed below, **Table 1b** and **c**). For details of the paired t-tests, see **Figs. S2, S3,** and **S4**. The differences in quantitation (Table S1) mean that some elements appear to be present as higher concentrations when using baseline subtraction (e.g., phosphorous, sulfur, chlorine, manganese) than when using baseline subtraction. The last of these is especially notable; manganese goes from apparently present at biologically relevant concentrations (baseline subtraction) to absent within uncertainty (blank spectrum correction).

**Figure 1:**
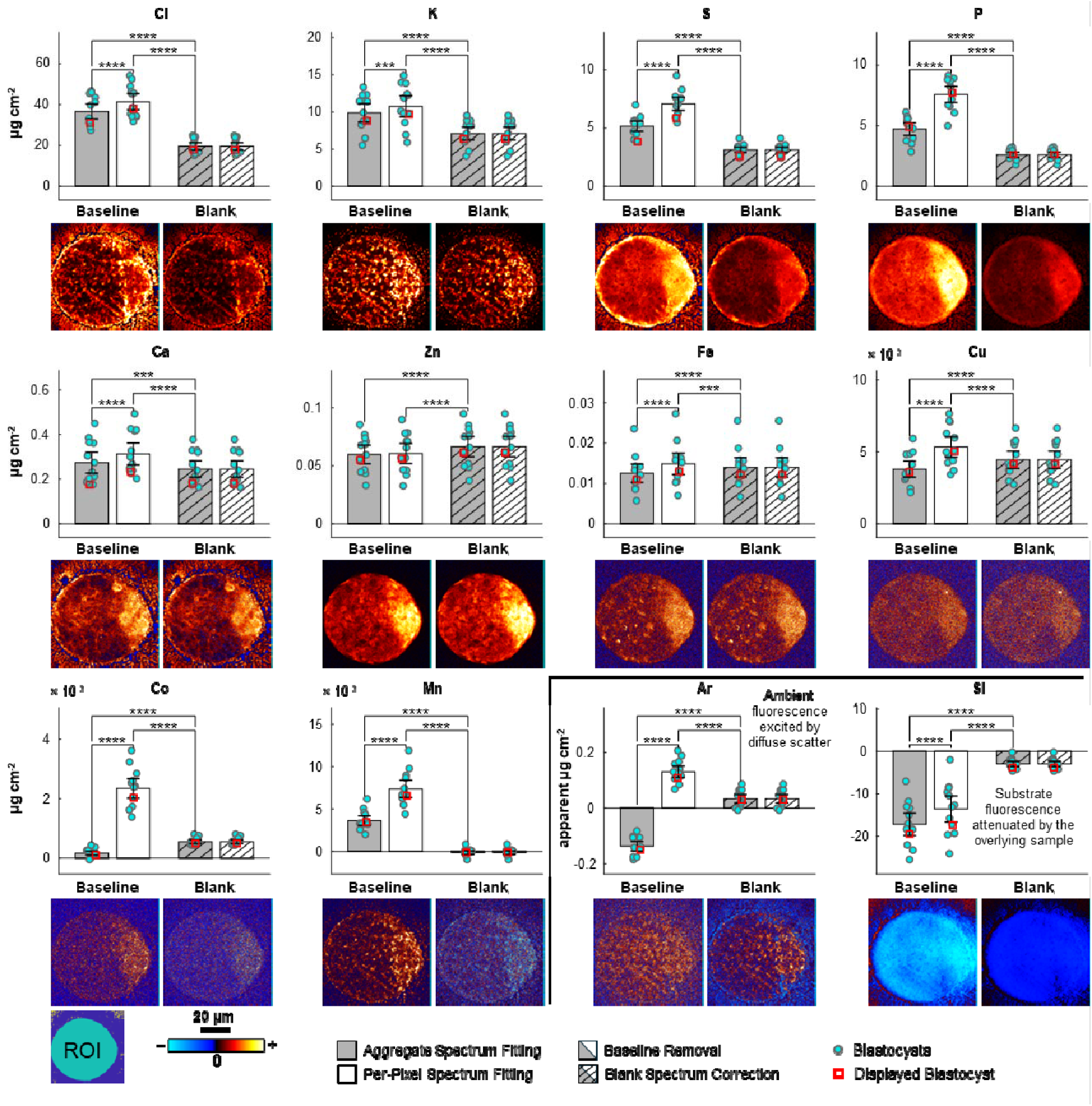
Differences in quantitative analysis between baseline and blank spectrum correction of per-pixel and aggregate spectra are biologically relevant. When analyzing the concentration of relevant elements across a set of 12 blastocytes, significant quantitative differences exist depending on whether baseline or blank background subtraction methods were used and whether analysis was performed on individual pixel spectra or the aggregate spectrum of image ROIs. Quantitatively, blank spectrum correction fitting of per-pixel and aggregate spectra differed negligibly with relative root-mean-squared-errors ca. 1×10^-14^ indicating only differences from computation precision. However, baseline subtraction of per-pixel and aggregate spectra never agreed, nor did either agree with quantitation from blank spectrum correction fitting. Bar plots correspond to the population mean. Errors bars are set at one standard deviation representing biological variation. Silicon is negative because it corresponds to the attenuated signal from the substrate. Each set of images are centered at zero intensity and scaled such that the magnitudes of the negative and positive extremes correspond to the greatest magnitude of the 1% and 99% quantiles. Statistical differences were determined using a paired-sample t-test: * p<0.05; ** p<0.01; *** p<0.001; **** p<0.0001.

**Table 1.** Relative percentage differences in quantification and population distribution from ROI analysis of the 12 blastocysts shown in Fig. 1, and tabulated in Table S1, resulting from baseline subtraction and blank spectrum correction of per-pixel and aggregate spectra. The comparison of blank spectrum correction for aggregate and per-pixel spectra is not shown because the answers were identical within computational precision (please refer to Table S2). Statistical differences were determined using a paired-sample t-test, with the results of unpaired-sample t-test shown as a courtesy.

| (a) | Baseline (Aggregate vs. Per-Pixel) |  |  |  | (b) | Aggregate (Baseline vs. Blank) |  |  |  | (c) | Per-Pixel (Baseline vs. Blank) |  |  |  |
| --- | --- | --- | --- | --- | --- | --- | --- | --- | --- | --- | --- | --- | --- | --- |
| | $\Delta$ Conc. | $\Delta$ Pop. Dist. | Paired T-Test | Unpaired T-Test | | $\Delta$ Conc. | $\Delta$ Pop. Dist. | Paired T-Test | Unpaired T-Test | | $\Delta$ Conc. | $\Delta$ Pop. Dist. | Paired T-Test | Unpaired T-Test |
| Si | 27% | -14% | $7.4 \times 10^{-7}$ | ns | Si | 470% | 327% | $1.0 \times 10^{-7}$ | $7.0 \times 10^{-9}$ | Si | 350% | 396% | $1.3 \times 10^{-5}$ | $6.8 \times 10^{-6}$ |
| P | -38% | -21% | $8.1 \times 10^{-11}$ | $8.0 \times 10^{-6}$ | P | 91% | 146% | $7.2 \times 10^{-8}$ | $5.0 \times 10^{-7}$ | P | 206% | 216% | $7.9 \times 10^{-10}$ | $1.3 \times 10^{-11}$ |
| S | -27% | -23% | $6.8 \times 10^{-10}$ | $1.2 \times 10^{-4}$ | S | 73% | 97% | $2.5 \times 10^{-9}$ | $8.6 \times 10^{-9}$ | S | 137% | 156% | $4.5 \times 10^{-10}$ | $5.4 \times 10^{-11}$ |
| Cl | -12% | -6% | $4.2 \times 10^{-6}$ | ns | Cl | 87% | 109% | $1.1 \times 10^{-8}$ | $3.3 \times 10^{-7}$ | Cl | 111% | 123% | $3.3 \times 10^{-9}$ | $1.3 \times 10^{-8}$ |
| Ar | -205% | -16% | $2.8 \times 10^{-10}$ | $4.5 \times 10^{-14}$ | Ar | -436% | 44% | $1.8 \times 10^{-8}$ | $7.0 \times 10^{-13}$ | Ar | 220% | 72% | $3.3 \times 10^{-8}$ | $1.3 \times 10^{-6}$ |
| K | -8% | -13% | $1.4 \times 10^{-4}$ | ns | K | 40% | 42% | $3.9 \times 10^{-8}$ | $3.7 \times 10^{-3}$ | K | 52% | 64% | $2.5 \times 10^{-7}$ | $8.8 \times 10^{-4}$ |
| Ca | -13% | -4% | $8.6 \times 10^{-7}$ | ns | Ca | 13% | 27% | $2.9 \times 10^{-4}$ | ns | Ca | 30% | 33% | $8.5 \times 10^{-7}$ | ns |
| Mn | -51% | -44% | $4.1 \times 10^{-8}$ | $3.5 \times 10^{-5}$ | Mn | 1902% | 182% | $9.6 \times 10^{-8}$ | $2.5 \times 10^{-10}$ | Mn | 3959% | 404% | $4.6 \times 10^{-8}$ | $1.7 \times 10^{-11}$ |
| Fe | -16% | -14% | $2.0 \times 10^{-6}$ | ns | Fe | -10% | -5% | $9.5 \times 10^{-9}$ | ns | Fe | 7% | 10% | $3.7 \times 10^{-4}$ | ns |
| Co | -93% | -79% | $3.2 \times 10^{-8}$ | $1.0 \times 10^{-7}$ | Co | -68% | 36% | $1.5 \times 10^{-7}$ | $7.3 \times 10^{-7}$ | Co | 365% | 543% | $2.2 \times 10^{-7}$ | $2.7 \times 10^{-9}$ |
| Cu | -29% | -21% | $3.1 \times 10^{-8}$ | $7.4 \times 10^{-3}$ | Cu | -13% | -6% | $8.5 \times 10^{-9}$ | ns | Cu | 23% | 19% | $4.8 \times 10^{-7}$ | ns |
| Zn | -1% | -7% | ns | ns | Zn | -10% | -9% | $1.4 \times 10^{-8}$ | ns | Zn | -9% | -2% | $8.1 \times 10^{-7}$ | ns |

Other elements (iron, cobalt, copper, and zinc) are underestimated by as much as a factor of 3 when using baseline subtraction. Baseline subtraction increases the apparent attenuation of the silicon fluorescence from the substrate and gives non-physical negative fluorescence from the ambient argon. In addition, the apparent variance, which reflects the biological population distribution, is generally larger when using baseline subtraction.

### Quantitative estimates using baseline subtraction are systematically dependent on experimental dwell time

Based on the comparison above, we hypothesized that uncertainty in the data could disproportionately affect the baseline-subtracted data, resulting in different quantitation depending on the noise level of the data. To test this, part of a murine ovarian follicle was imaged four times at the BNP (Bionanoprobe) with a 180 nm step size, using dwell times of 5 ms, 10 ms, 50 ms, and 1000 ms. The areal densities for calcium and copper of the scanned region of the follicle were calculated using either blank spectrum correction or baseline subtraction prior to fitting and are presented for the different dwell times along with the elemental maps (**Fig. 2**). The same pixels used for blank spectrum correction were applied to baseline-removed data when adjusting each element by its background mean. Blanks from each dwell time collection (**Fig. S1**) remain consistent, aside from signal-to-noise (S/N) variations, as shown in previous work,^27, 42^ and recapitulated here.

**Figure 2:**
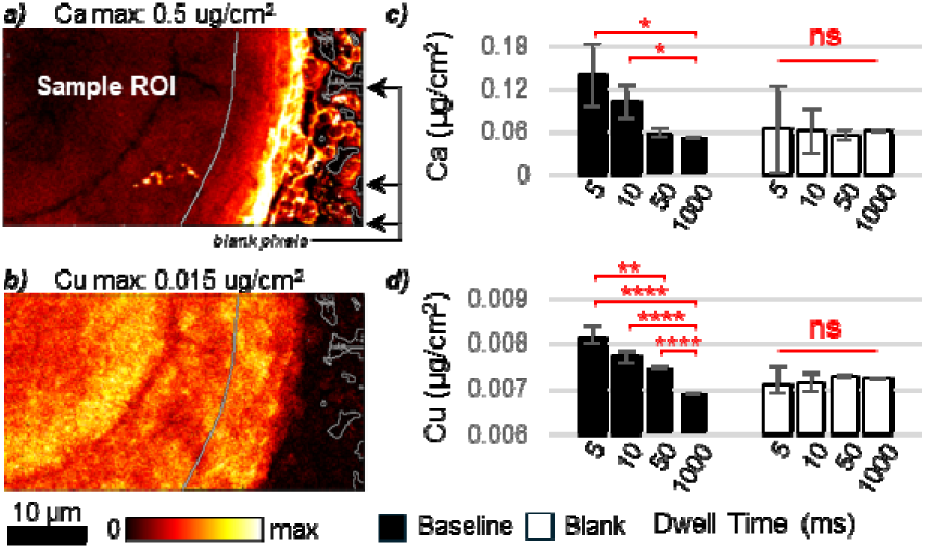
Varying experimental dwell time changes quantitative estimates for baseline subtraction but not for blank spectrum correction. Elemental concentrations in a tissue section of a murine ovarian secondary follicle were measured at four different per-pixel exposure times and analyzed using two different x-ray fluorescence analysis methods. Exposure time should not change calculated concentrations; that is the case with blank spectrum correction, but not with baseline subtraction for background signals. This can be seen by looking at the XFM maps for calcium (a) and copper (b) of the murine ovarian secondary follicle and comparing the mean concentration of each sample for calcium (c) and copper (d) for the two analysis methods and four per-pixel exposure times. Errors bars are set at one standard deviation representing biological per-pixel variation. Since each dwell time measurement consisted of ca 35000 pixels, statistical differences were determined using a two-sample z-test: * p<0.05; ** p<0.01; *** p<0.001; **** p<0.0001. There were no statistically significant differences between the blank spectrum

When comparing the fitted calcium and copper concentrations across the four dwell times, none of the quantitative estimates from baseline subtraction agreed. The apparent concentration of both calcium and copper decreases monotonically with increasing dwell time when using baseline subtraction. In contrast, results from blank spectrum correction were independent of dwell time.

While the elemental content is expected to vary between replicate samples, a single sample measured at different dwell times should not change aside from changes in S/N.

### Inconsistencies in baseline removal lead to potential errors in quantitation

Since the total elemental signal in an image (e.g., the 1×1 mm square of the murine ovarian follicle in **Fig. 3**) must be independent of how post collection spectra are binned, adjustments in bin size can be used as a surrogate for mimicking the effect of dwell time, but with the benefit of having an absolute quantity available for comparison. Therefore, we took a single image of an ovarian follicle and performed NxN binning of the spectra across the xy plane. Doing this held the data content constant while increasing the counting statistics (at the cost of decreased spatial resolution); this allowed us to test the relationship between the S/N ratio and apparent quantitation, independent of any changes in the measurements or instrumental sensitivity. This NxN binning is similar to what is often used for standard reference measurements and biologically relevant image ROIs when integrating the relevant pixel spectra to a single spectrum prior to fitting,^45–47^ and is analogous to separating an image into equally spaced image ROIs.

**Figure 3:**
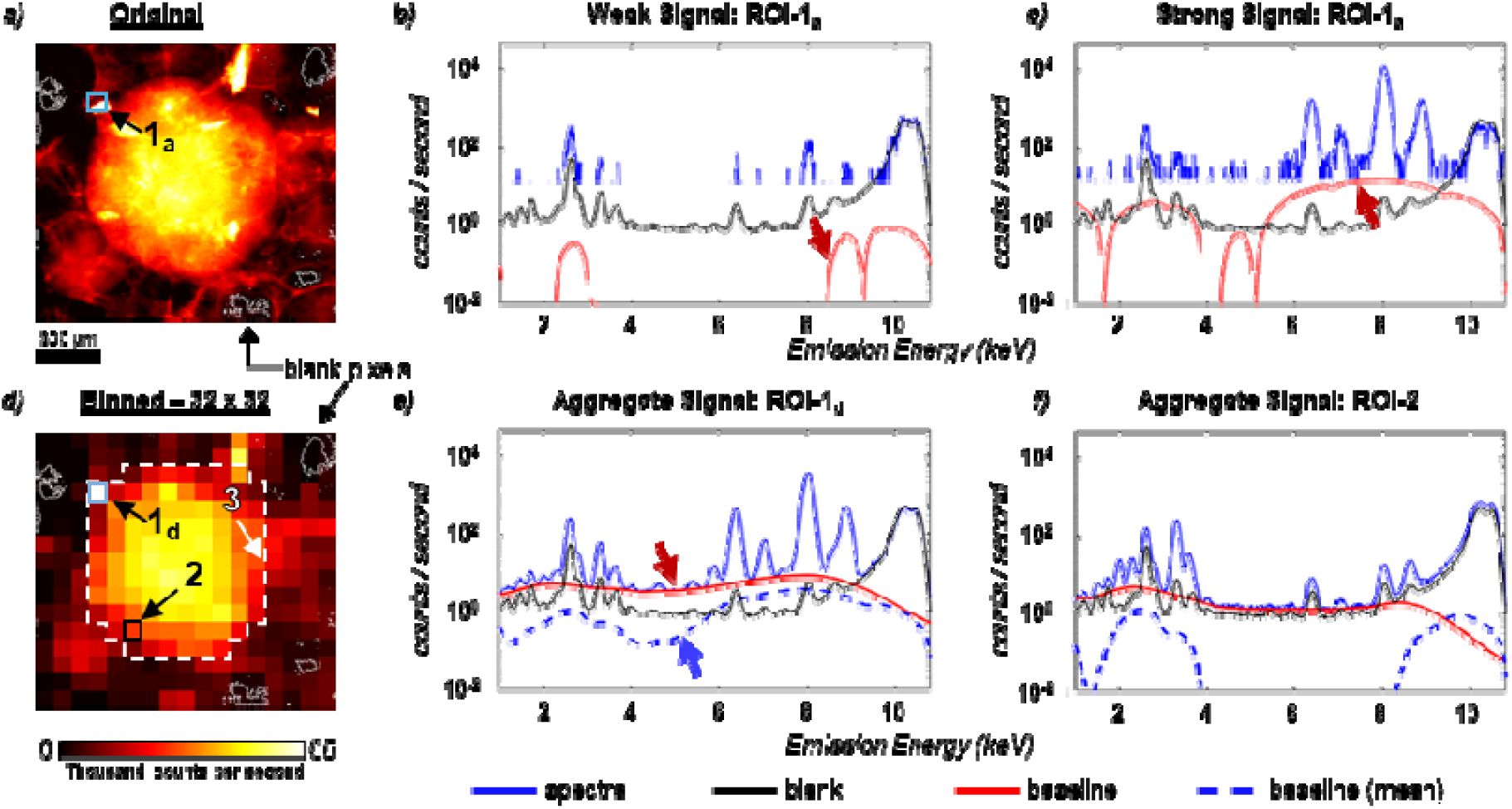
Baseline amplitude and morphology are dependent on total dwell time resulting in variable background correction of XFM data when using baselines, but not when using blanks. Using the integrated XFM spectra image (a) for a murine ovarian secondary follicle, a weak (b) and strong signal (c) were selected from ROI-1_a_. For both signals, it is especially difficult to correctly model a background. Significantly increasing the counting statistics at each pixel by 32×32 pixel binning of the spectra (d) insufficiently ameliorates this baseline variation as seen in the aggregate signals of the 32×32 binned pixels in ROI-1_d_ (e) and ROI-2 (f). Although the baselines benefit from greater statistical significance, they remain inconsistent. The spectra from the energy dispersive detector are shown in blue, the baseline in red, the blank from the indicated pixels in black, and (for the integrated spectra in e and f) the average baselines from the original unbinned data shown in (a). The high aggregate statistics of the blank spectrum correction method avoid the artifacts caused by inaccurate estimation of backgrounds by baselines. ROI 3 corresponds to the region used to calculate the aggregate signal from the whole follicle and is analogous to the longest simulated dwell time in Figure 6.

The raw (unbinned) image of the murine ovarian follicle is shown in **Fig. 3a**, with ROI-1_a_ corresponding to a 32×32 neighborhood of pixels. From ROI-1_a_, representative spectra of a weak signal (**Fig. 3b**) and strong signal (**Fig. 3c**) are shown along with the calculated baselines and empirically measured blank. For the weak signal, the baseline (red line) underestimates the continuum, while overestimating it for most of the strong signal.

It was possible that variations in baseline amplitude and morphology could be mitigated with a sufficiently well-defined signal. To test this, spectra from two aggregate regions from the 32×32 binned data (**Fig. 3d**) are compared (**Fig. 3e** and **f**). **Fig. 3e** corresponds to the spectrum from ROI-1_d_ and is identical to the aggregate signal of the 1024 spectra from ROI-1_a_. Where the baseline in ROI-1_d_ significantly overestimates the continuum relative to the blank (black line), the baseline for the aggregate spectrum in ROI-2_d_ (**Fig. 3f**) only slightly overestimates the continuum. Still, baseline removal remains inconsistent, and neither baseline from ROI-1_d_ or ROI-2_d_ agree with their associated aggregate baseline from the raw data (dashed blue lines). In both cases, the baselines from the aggregate spectrum had a greater amplitude than the aggregate of the baselines from the individual pixel spectra. These data imply potential systematic differences from baseline removal.

### Aggregating spectra before baseline removal recapitulates the quantitative shifts seen with increasing dwell time

Following the NxN binning, the resulting spectra were baseline corrected and then integrated (**Fig. 4**). As the per-pixel counting statistics of spectra increase, the amplitudes of the baselines also increase resulting in the apparent decreases in the area under the elemental peaks of interest. This explains the decrease in the amplitudes of the integrated spectra following baseline removal. Qualitatively, the dependency of baseline removal on ROI size or dwell time is best seen by the ca. 10-fold decrease in spectral amplitude between 4 keV and 8 keV. Quantitatively, however, every peak is affected. Comparing the peak amplitudes from the original spectra to those with 256×256 bins, the apparent amplitudes of phosphorus, potassium, calcium, iron, copper, and zinc all decreased by 30-40%, with manganese decreasing by 80% (**Fig. 4, inset panels**). Since the original dwell time was 0.05 seconds, the effective dwell time for a NxN binned dataset is 0.05×N^2^, giving effective dwell times of over 3000 seconds for the largest bins.

**Figure 4:**
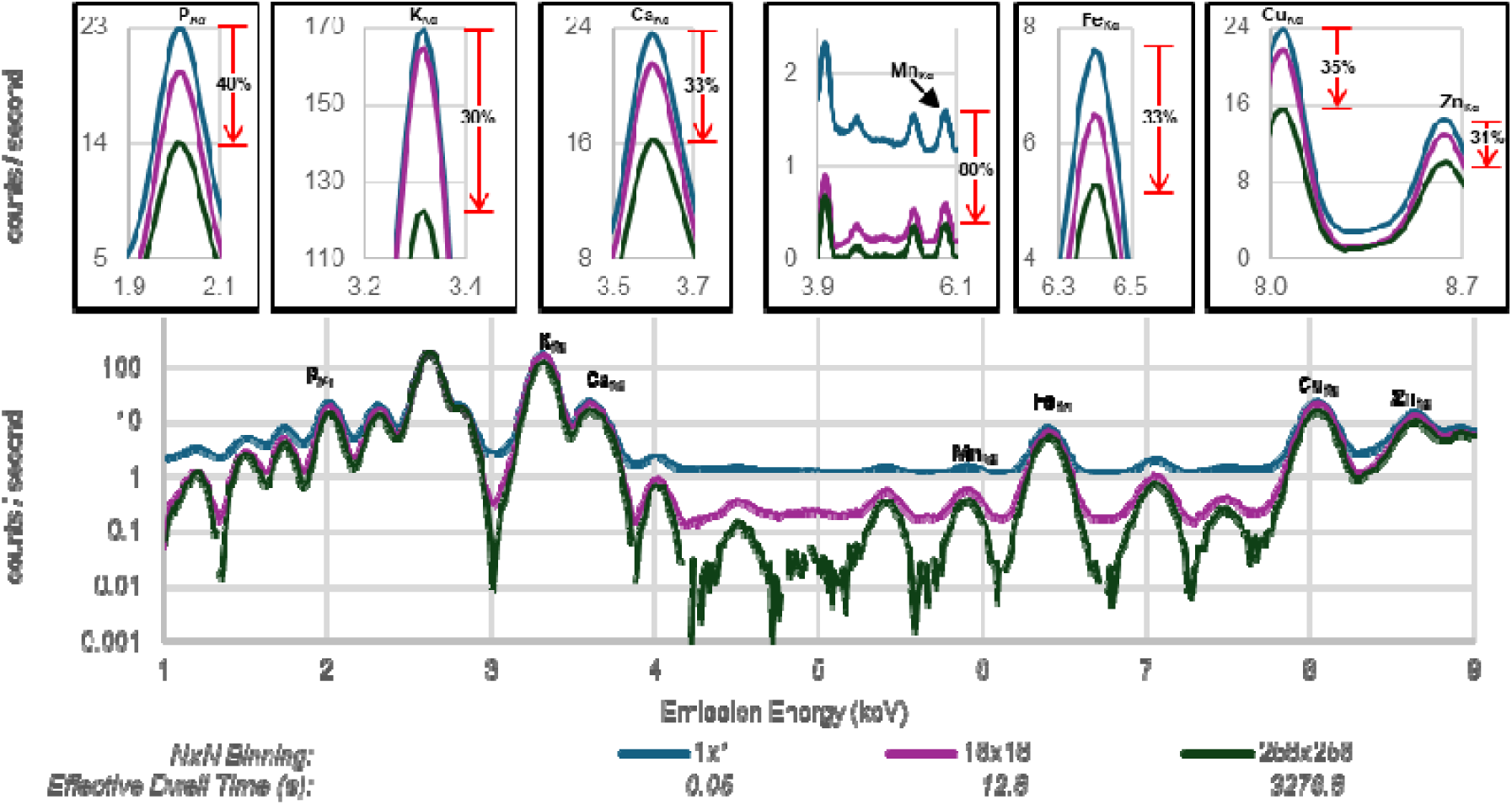
Using baseline subtraction with increasing dwell time results in systematic decrease of elemental peak areas. This is demonstrated using pixel binning of X-ray fluorescence spectra data of the murine ovarian secondary follicle as shown in Fig. 3 by 16×16 and 256×256 binning. The resultant spectra from each of the three groups were baseline corrected and then integrated and plotted in units of counts per second. As the per-pixel statistics increases with the NxN binning, there is a corresponding systematic decrease in each element’s peak intensity, with ca. 35% decreases for silicon, potassium, calcium, iron, copper, and zinc, and an 80% decrease for manganese.

Each NxN binned dataset was fitted using both blank spectrum correction and baseline removal, and the values were left in units of counts/second. To avoid any differences in quantitation that may arise from the use of different software packages, both baseline removal and blank spectrum correction were performed using M-BLANK with identical parameters to ensure consistent quantitative comparisons. The baseline corrected and blank-corrected results were compared by taking the ratio of the former to the latter (**Fig. 5a**). As demonstrated above, results from blank-corrected data are independent of dwell time (**Fig. 2c** and **d**), thus changes in the ratio are due to changes in baseline corrected data. This is further demonstrated in **Fig. 5b** and **c**, which show the ratios of the total fitted image content from the NxN binned datasets from both analyses ratioed to the total fitted content of the unbinned image. Note that blank-corrected results vary negligibly from unity, while baseline corrected results systematically decrease. Consequently, one can use blank spectrum corrected data as an absolute reference. From there, a quantitative evaluation can be made of any potential error caused by baseline subtraction.

**Figure 5:**
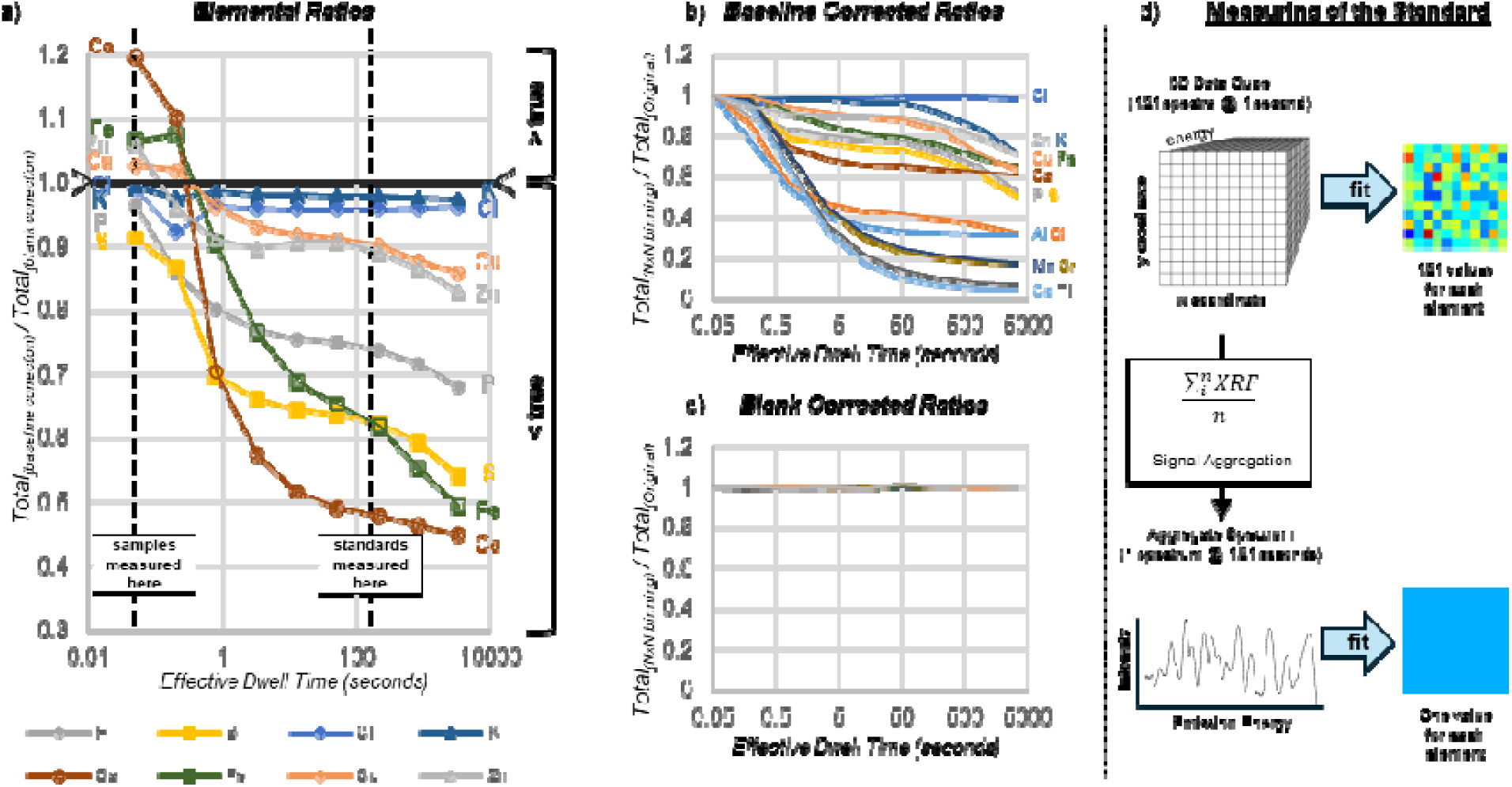
Estimation errors from baseline subtraction can be significantly compounded when mass calibration is performed by comparison to standards measured with significantly different counting statistics. (a) Using the total quantitation from the NxN binned XFM data from baseline subtraction and blank spectrum correction, the former was ratioed to the latter identifying a systematic change from overestimation to underestimation. To show this systematic error arises solely from baseline subtraction, the quantitative stability for baseline subtraction (b) and blank spectrum correction (c) are shown by ratioing the total fitted image content from each NxN binned dataset to the total content from the original data. The ratios from baseline subtraction (b) vary wildly, while the ratios from blank spectrum correction (c) remain constant at unity. (d) The copper image for the mass calibration standard imaged with 11 pixels on edge and collected at a 1 s dwell time, was subsequently integrated to a pseudo single-point spectrum prior to fitting. However, this can lead to a compounding of errors which can be understood by calibrating the calcium signal in (a) at 50 ms against itself at 120 s. The 20% overestimation at 50 ms and 50% underestimation at 120 s compound the error at both ends leading to a 140% overestimation of calcium at 50 ms.

### Mass calibration compounds error introduced by baseline subtraction

As the effective dwell time increases, the baseline-corrected quantitation for calcium, iron, copper, and zinc changes from being overestimated to being underestimated (**Fig 5a**). This trend of overestimation (at low counts) to underestimation (at high counts) is important when considering how XFM mapping data is typically collected and then mass calibrated. Samples are typically collected with very short dwell times (e.g., the dashed vertical black line on the left at 50 ms in **Fig. 5a**) to maximize sampling frequency; whereas, standards are typically collected with very long dwell times (e.g., the dashed vertical black line on the right at 120 seconds in **Fig. 5a**) or are aggregated from multiple pixels of a standard into a high-count pseudo single point spectrum (**Fig. 5d**) to minimize uncertainty of the calibration measurement. The quantitative impact of this can be understood by considering the ratios of the total elemental content of the murine ovarian secondary follicle (**Fig. 5a**) and calibrating it against itself. For calcium, the apparent counts at 50 ms are 20% overestimated, while the apparent counts at 120 s (analogous to a high counting statistics measurement of a standard reference material) are 50% underestimated. For a sample measured with 50 ms dwell and a calibration measured 120 s dwell, these effects would combine to overestimate Ca by 2.40-fold.

### Baseline subtraction estimates depend on the degree of peak separation and peak signal-to-background ratios

Here, we define signal-to-background (S/B) as the spectral contrast, or peak intensity relative to the local baseline continuum. To explore how variable peak separation and S/B ratios influence baseline variations and affect quantitative accuracy, we constructed a simple model consisting of two Gaussian functions riding on a constant zero-order linear background and explored the apparent peak area as a function of dwell time dependent statistical noise, calculated from a Poisson distribution. Using this model, simple spectra were calculated for theoretical dwell times between 50 ms and 5000 s. For an image with 50 ms dwell time, this corresponds to image ROI sizes between 1 pixel (as in the blue spectra in Fig. 3b and c) and 100,000 pixels (as in ROI-3d in **Fig. 3**). For long dwell times or large ROIs (**Fig. 6 c**, and **h**), baseline subtraction generally overestimated the background while underestimating it at shorter dwell times (**Fig. 6 a**, and **f**), as seen in our experimental data. By varying the separation width between the two peaks the amount of overestimation and underestimation changed. This can be seen by comparing **Fig. 6b** and **g** which only differ by the separation width between the two peaks. With a peak separation of 3 standard deviations (**Fig. 6b**), baseline overestimation resulted in underestimation of both peak areas (**Fig. 6d**). However, when the same peaks were separated by 6 standard deviations (**Fig. 6f**), the baseline almost correctly predicted the background for peak 1 but underestimated the background for peak 2. Due to the opposite sign of the high-noise errors (overestimation) and incomplete-separation errors (underestimation), better separated peaks (Fig. 6i) give slightly worse estimates at short dwell times (Fig. 6d). The same simulation for blank spectrum correction was consistent with previous findings with < ±1% error at 50 ms and < ±0.1% error for the longest dwell times (**Fig. 6 e** and **j**). Lastly, S/B (different from S/N) governs susceptibility to baseline bias: as the peak’s contrast against the continuum decreases, small changes in the estimated baseline consume a larger fraction of the true peak area, amplifying the over/underestimation (Fig. 6a–d, f–i). This distinction (S/N vs S/B) explains why under baseline subtraction the peak with S/B ≈ 0.25 is perturbed more than the peak with S/B ≈ 1. In contrast, blank spectrum correction preserves a true zero reference and the peak–continuum contrast, so its errors remain near zero across S/B (Fig. 6e and j).

**Figure 6:**
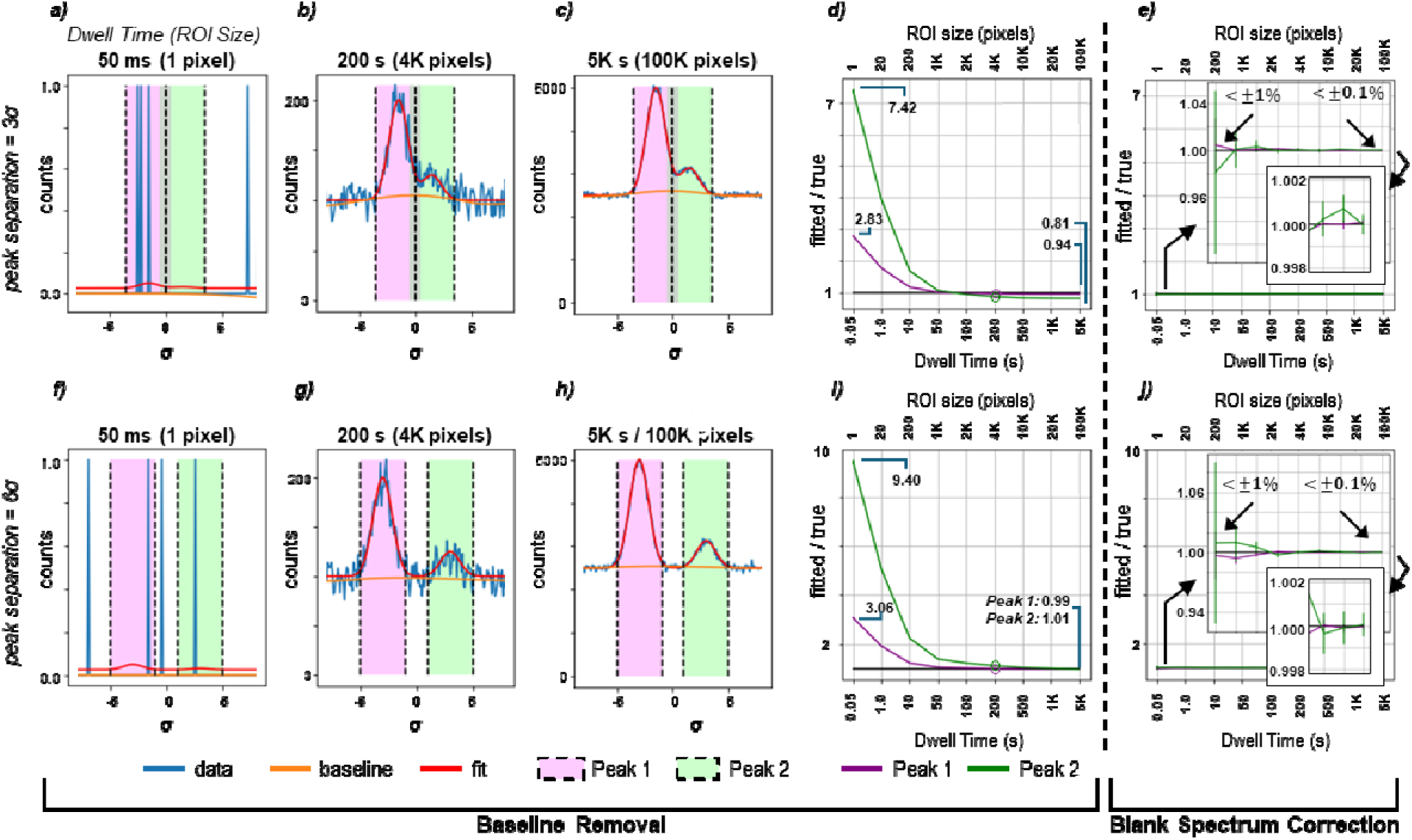
Baseline distortion is also systematically dependent on peak separation and the signal-to-background intensity ratio. We used a simple model consisting of two Gaussian functions riding on a constant linear background to explore the apparent peak area as a function of statistical noise, calculated from a Poisson distribution. The model was defined to have a signal-to-noise ratio of 1 for 1 second. We started with a single Gaussian with peak height of 1 and a background of 1. To study the effect of peak overlap, a second, smaller peak (peak height = 0.25) was added. Both Gaussians had a full-width half-maximum of 1 and were separated by 3 (a-e) and 6 (f-j) standard deviation units. Simulations were done for dwell times ranging from 50 milliseconds to 5,000 seconds (effective image ROI sizes of 1 pixel to 100,000 pixels), with representative simulations for 50 milliseconds (a and f), 200 seconds (b and g), and 5,000 seconds (c and h) shown. For each of the resulting data sets, baseline subtraction was modeled as a SNIP baseline calculated using the pybaselines implementation of the SNIP algorithm (https://pypi.org/project/pybaselines/) with 5 point moving-average smooth (keyword smooth_half_window = 2) and a snip-width set at the full-width-at-half-max (keyword max_half_window = σ). Blank spectrum correction was simulated by adding Poisson based noise corresponding to a 500 second measurement to the linear background. For each simulation, the ratio of fitted counts to the true counts was determined. In order to estimate the reliability of the simulations, each condition was repeated 10,000 times and the resulting ratios were averaged to give the mean and standard deviation.

## Discussion

Blank spectrum correction of XFM data yields improved accuracy in quantitation as compared to baseline subtraction. The latter can distort the interpretation of biological data. This can be seen from comparison of these two approaches applied to per-pixel and aggregate ROI spectra of the preimplantation blastocyst mouse embryos (**Fig. 1**, **Table 1**). Where blank spectrum correction yielded identical quantitative results, baseline subtraction never agreed, and both the quantitation and population distributions from fitting aggregate ROI spectra were always less than the same quantitation and population distributions from fitting per-pixel spectra. This is because baselines exhibit notable variations depending on the spectral quality (S/N).

Changing the pixel dwell time used during data collection or the post-collection pre-processing of data by spectral aggregation of image ROIs will result in different baselines and therefore different estimates of the elements of interest. At low measured signal counts, baselines underestimate the background leading to an overestimate of the sample (e.g., with shorter dwell times or individual pixel spectra), while at high measured signal counts the baseline is overestimated and the element counts underestimated (e.g., with longer dwell times or aggregate ROI pixel spectra). These variations are additionally dependent on the relative signal-to-background ratio, and the closer to zero an element intensity is, the more important true zero becomes. As such, the lower the spectral contrast (S/B) the greater the magnitude of the baseline bias on the peak areas. Baseline variations also depend on the degree of peak overlap which will further complicate analyses with overlapping signals (e.g., iron and manganese in the presence and absence of gadolinium).

Although our study used results for preimplantation mouse blastocysts, we also compared results from early-stage blastocysts, as well as 8-cell and morula stage embryos (**S5, S6, S7**). The developmental biology implications of these data sets are discussed elsewhere.^48^ Statistically significant differences in elemental measurements and population distributions were identified in the blastocyst samples. The differences between values obtained from baseline subtraction and blank spectrum correction in the cases of iron, copper and zinc are within experimental uncertainties arising from biological variation in these data sets; however, we note that such differences will always be sample and beamline dependent. For phosphorus through calcium, they deviate largely and are also particularly pronounced for manganese and cobalt. Baseline subtraction caused a manganese overestimation of ∼19-fold for aggregate fitting or ∼40-fold per-pixel fitting. For cobalt, baseline subtraction led to significant underestimate (∼70% for aggregate fitting but a > 3-fold overestimate for per-pixel fitting. Errors of this magnitude can have important implications for biology, leading to misinterpretation as manganese intoxication, cobalt intoxication or cobalt deficiency. Any of these findings would be notable as both elements are known to influence reproductive health: manganese plays roles in hormone regulation, oocyte maturation and reproductive function, whereas cobalt influences trophoblast proliferation and embryo development.^49^

Since baseline subtraction yields quantitation that is dependent upon the amount of signal aggregation, comparisons between ROIs of different sizes can be problematic. The systematic shift from overestimation to underestimation with increasing ROI size introduces inconsistencies in quantitative analysis making it difficult to draw accurate conclusions. By extension, since changes in signal intensity result in variable baselines, this confounds comparison between data collected at different beamlines or different synchrotrons.

Limitations of the blank spectrum correction approach include the assumption that background contributions are uniform across the sample. This is a reasonable assumption for synchrotron-based XFM analysis of thin samples on uniform substrates, or thicker samples mounted to the backside of x-ray transparent film; however, in methods such as PIXE^50^ or benchtop XRF where Bremsstrahlung contributions vary across the sample, more complicated methods of background correction may be required. More broadly, if the background is spatially non-uniform or spectrally mismatched between the chosen off-sample region and the ROI, any off-sample–based correction is compromised: spectrum-level blank subtraction and post-fit “background subtraction” (after baseline subtraction) will both inherit the same bias. The remedy is practical rather than conceptual—select representative blank ROIs, verify background uniformity (or, if necessary, stratify blanks by region or model the background explicitly)—not an inherent limitation of spectrum-level blank correction.

In conclusion, blank spectrum correction was demonstrated to be more quantitatively consistent when compared to baseline subtraction. We identified three factors that influence the extent to which blank spectrum correction outperforms baseline subtraction: inherent S/N (**Figs. 2-5**), inherent signal-to-background (**Fig. 6**), and degree of peak overlap (**Fig. 6**). These factors are present in fitting of both the reference and the sample spectra and if these are measured under different S/N conditions or treated differently (as is the case when aggregating the standard measurement to a high-count pseudo single point spectrum), errors arising from baseline subtraction can be compounded, and the resulting apparent concentrations can be significantly overestimated or underestimated. Given that X-ray spectra present signal intensities across orders of magnitude with variable degrees of peak overlap, variable S/N, and variable signal-to-background, we support the use of blank spectrum correction in analyzing XRF spectra. Although we have focused on interpretation of XRF data, we expect similar problems in related spectroscopic techniques that use baseline subtraction. Such discrepancies could potentially explain, in part, the nonlinear nature of calibration curves in laser ablation imaging and the putative need for weighted least squares.^51^

## Methods

### Sample Preparation

CD-1 mice were purchased from Envigo (IN, USA) and bred in-house at the Northwestern University Center for Comparative Medicine animal facility. Animals were given food (Teklad 2020X, Envigo) and water *ad libitum*, and treated in accordance with the National Institutes of Health Guide for the Care and Use of Laboratory Animals. Mice were kept in a 14-hour light, 10-hour dark cycle with constant temperature and humidity. All protocols were approved by the Northwestern University Institutional Animal Care and Use Committee.

Preimplantation embryos were isolated from CD-1 female mice (6-8 weeks old) after hyperstimulation with 10 IU intraperitoneal pregnant mare’s serum gonadotropin (PMSG, EMD Bioscience, San Diego, CA, USA). After 44-46 hours, mice were superovulated with 10 IU intraperitoneal human chorionic gonadotropin (hCG, Sigma-Aldrich, St. Louis, MO, USA). Two female mice were paired with one CD-1 male mouse (10-12 weeks old) for trio breeding. Blastocyst embryos were collected from the oviducts and uterine horns at 120 hours post-hCG. Oviducts and uteri were collected into dissection media containing Leibovitz’s L-15 medium (L-15, Sigma Aldrich, St. Louis, MO, USA) supplemented with 0.1% polyvinyl alcohol (PVA, Sigma Aldrich, St. Louis, MO, USA). The tissue was cut 8-10 times with scissors to release the embryos. Embryos were collected and washed in potassium complex optimized media (KSOM, EMD Millipore, Burlington, MA, USA) that was pre-equilibrated in an incubator (37⍰C, 5% CO_2_ in air). Embryos were prepared for x-ray fluorescence microscopy by washing in 100 mM, pH 7 ammonium acetate to remove metal salts. Next embryos were loaded on a 1.5 mm x 1.5 mm by 500 nm thick silicon nitride window (Norcada Inc., Canada), air dried and stored at room temperature until mapping.

Ovarian follicles with two layers of somatic cells (secondary follicles) were isolated from postnatal day 12 female CD-1 mice via mechanical digestion as described elsewhere.^43^ Briefly, ovaries were dissected out of the ovarian bursa and mechanically disrupted via ‘flicking’ with insulin gauge syringe needles (ThermoFisher, MA, USA) in Leibovitz’s L-15 medium (ThermoFisher, MA, USA) supplemented with 1 mg/mL polyvinyl alcohol (MilliporeSigma, MA, USA) until their follicles were released. Individual follicles were collected and washed in 100 mM, pH 7 ammonium acetate solution (MilliporeSigma, MO, USA) supplemented with 1 mg/mL PVA to remove metal salts in the media before being placed on an intact 1.5 mm x 1.5 mm by 500 nm thick silicon nitride window (Norcada Inc., Canada). Excess liquid was removed by wicking with a mouth pipette. The windows were placed on a 37°C heated stage for less than one minute until dry and stored at room temperature until mapping.

### Synchrotron X-ray Fluorescence Microscopy

Using beamline 2-ID-E of the APS, X-ray fluorescence spectra of 12 blastocysts were collected in fly scan mode at 300 nm step sizes and 50 ms dwell times with incident energy of 10 keV selected using a Kohzu-Si(111) double crystal monochromator from a 3.3 cm undulator source with a ΔE/E of 1.4×10^-4^ at 12 keV. Samples were oriented at 15° to the incident X-ray beam with the detector positioned at 90° to the incident X-ray beam to minimize the intensity of Compton (or inelastic) scattered X-rays. The beam spot was focused to a full-width at half-maximum (FWHM) of 300 nm using a Fresnel zone plate.^52^ Fluorescence was measured using a Vortex 4-element silicon drift detector running Digital X-ray Processor electronics from X-ray Instrumentation Associates, and normalized to a nitrogen filled upstream ion chamber attached to a Stanford Research SR570 low noise current to voltage preamplifier. The XFM images for the 12 blastocyst samples were analyzed using both blank and baseline subtraction prior to spectral deconvolution.

Using the BNP^53^ at beamline 9-ID of the APS, X-ray fluorescence spectra of ovarian secondary follicles were collected in fly scan mode at various step sizes and dwell times with incident energy of 10 keV selected using a double crystal monochromator. Samples were oriented at 15° to the incident X-ray beam with the detector positioned at 90° to the incident X-ray beam to minimize the intensity of Compton scattered X-rays. The beam spot was focused to a FWHM of 100 nm using a Fresnel zone plate,^52^ and fluorescence was measured using a Vortex 7-element detector running Xspress3 electronics by Quantum Detectors. Fluorescence counts were normalized to the beam intensity as measured using a downstream photodiode.

### Spectral Fitting

All XRF data in this work have been fitted either as individual pixels or as the aggregate spectrum from user chosen image ROIs. For aggregate spectrum fitting, all pixels corresponding to an image ROI, e.g., the sample or background, are summed into a single spectrum prior to baseline subtraction or blank spectrum correction (see below). Fitting of XRF data (collected below) was performed using both the M-BLANK^27^ and MAPS v9.0^32^ software packages, with blank spectrum correction being performed in M-BLANK^27^ and baseline subtraction performed in MAPS^32^, unless otherwise stated. Notably, there are many programs that use a baseline subtraction approach^26–32, 36–41^; MAPS was used because it was readily available. Peak shape definitions and specifics of fitting can be found in the supplemental information associated with the article by Crawford et al.^27^ Briefly, fluorescence emissions were modeled as modified Gaussians as outlined in the *Handbook of X-ray Spectrometry*^54^ and by Phillips and Marlow ^55^, with a slight correction by Campbell and Maxwell^56^. Emission energies, branching ratios^57–58^, and fluorescence yields were referenced using the xraylib^59^ program and the *CXRO X-ray Data Booklet*^60^, with branching ratios for M-BLANK^27^ empirically optimized^57^ during parameter creation. Once calculated, peak shapes were held constant and used in linear least squares fitting of the spectral data for M-BLANK^27^ and non-negative matrix factorization for MAPS.^32^ MAPS fitted data were taken directly from the original studies, using the baseline and fitting parameters optimized therein.^43, 48^ Fitted fluorescence values were converted to areal concentration (µg cm^-2^) by comparison to an Axo thin film standard (Ca, 1.931 µg cm^-2^; Fe,0.504 µg cm^-2^; Cu, 0.284 µg cm^-2^; Axo Dresden GmbH, Gasanstaltstr. 8b, 01237 Dresden, Germany).

### Baseline Subtraction and Blank Spectrum Correction of XRF

For XRF, the detected signal (F) contains fluorescence from the sample (S), an x-ray fluorescence continuum (C), and contributions from background fluorescence (B) from such things as the substrate (i.e., the Si_3_N_N_ wafer) and the instrument/hutch, such that F = S + C + B. Baseline subtraction of XRF accounts for C by calculating baseline (L), such that the residual F_SB_ = S + B can be calculated and then fit. After the data are fit, B is accounted for by selecting an off-sample area, taking the average for each fitted element, and then subtracting the mean value of that region from the fitted elemental map (or aggregate ROI value). Alternatively, blank spectrum correction of XRF accounts for both C and B by taking the mean spectrum of the same off-sample ROI, and then subtracting that spectrum from the spectra to be fitted. With baseline subtraction, it calculates L to model C. This assumes that L correctly models C such that C – L = 0. If not, then C – L = Δ, where Δ is the error in the baseline’s accounting of the continuum.^27^

### Dwell Time Experiments

One of the ovarian secondary follicles scanned at the BNP (Bionanoprobe, Advanced Photon Source, Argonne National Laboratory, USA) was imaged such that a region containing the oocyte, zona pellucida, and layers of somatic cells was positioned in the path of the X-ray beam. The sample was rastered at 180 nm steps along x and y. Identical scan settings were used to image the same region in triplicate with associated dwell times of 5 ms, 10 ms, 50 ms, and 1000 ms. These data were then processed and fitted using baseline correction in MAPS and blank spectrum correction in M-BLANK for comparison.

### Signal Aggregation via Pixel Binning

To explore the effect of signal aggregation, or region of interest (ROI) choice, on the fitted results, a 1 mm^2^ image of a murine ovarian secondary follicle was scanned at the BNP at a step size of 2 µm in both x and y. The 2 µm x 2 µm image pixels were then digitally combined using progressively larger NxN pixel binning (N=2, 4, 8, 16, 32, 64, 128, 256) as “image ROIs". The spectra of these “image ROIs” were then processed using baseline subtraction and blank spectrum correction and analyzed at the level of the spectra and post fitting. For baseline subtraction, the baselines for each dataset were calculated using the MAPS software package and then used in lieu of the blank to correct the spectra. All data reduction and fitting were performed using M-BLANK (with identical parameters) to avoid differences in quantitative estimates that could arise from the use of different software packages.

### Simulated Peak Spectra

To study the relationship between baseline artifacts, peak overlap, and variable signal-to-background ratios, we constructed a simple model consisting of two Gaussian functions riding on a constant zero-order linear background with statistical noise calculated from a Poisson distribution. The model was defined to have a signal/noise ratio of 1 for 1 second. The background and two Gaussians had relative amplitudes of 1, 1, and 0.25 respectively. Both Gaussians had a FWHM of 1 and were separated by either 3 or 6 standard deviation units. Simulations were done for effective dwell times ranging from 3 seconds to 10,000 seconds. For each of the resulting data sets, a SNIP baseline was calculated using the pybaselines implementation of the SNIP algorithm (https://pypi.org/project/pybaselines/) with 5 point moving-average smooth (keyword smooth_half_window = 2) and a snip-width set at the full-width-at-half-max (keyword max_half_window = σ); The blank was simulated by taking the linear background and adding Poisson noise simulated for a 500 second measurement. For each simulation, the ratio of baseline-subtracted area to the true area was determined. To estimate the reliability of the simulations, each condition was repeated 10,000 times, and the resulting ratios were averaged to give the mean and standard deviation. For thoroughness, to evaluate the effect of changing the snip-width in calculating the baseline, see Figure S8.

### Statistics

Statistical p-values of the elemental content for the murine ovarian blastocysts in Fig. 1 were calculated using a paired t-test: * p<0.05; ** p<0.01; *** p<0.001; **** p<0.0001. Statistical p-values of the elemental content for the dwell time comparisons of the murine ovarian secondary follicle in Fig. 2 were calculated using a Z-test with a Spearman’s rank test to check for a monotonic relationship between dwell time and quantitative estimates.: * p<0.05; ** p<0.01; *** p<0.001; **** p<0.0001.

## Supporting information

Table of Content Graphic

## AUTHOR INFORMATION

### Author Contributions

AMC wrote all software and algorithms. AMC, JEPH, CJ, and KWM wrote the manuscript with feedback from TKW and TVO. SG assisted in statistical analyses. JB provided the murine early-stage blastocysts, late-stage blastocysts, 8-cell, and morula stage embryos. YC provided the ovarian follicles. QJ assisted in XFM data collection at APS. All authors have given approval to the final version of the manuscript.

## Acknowledgements

We thank Vincent R. Ragusa (Department of Microbiology, Genetics, and Immunology, Michigan State University) for his helpful discussions and support. We thank Arthur Glowacki, Olga Antipova, Si Chen, Evan Maxey, and Barry Lai of the X-ray Microscopy Group at the Advanced Photon Source of Argonne National Laboratory.

