## Supplemental Information for "Blank Spectrum Correction as a Robust Solution to Artifacts in Quantitative X-ray Fluorescence Mapping"

ABSTRACT:

X-ray fluorescence microscopy (XFM) continues to develop as a powerful quantitative technique for high resolution, label-free, elemental mapping of biological, environmental, and material samples. Methods for rigorously fitting spectra, increasing throughput, accounting for background signals, and deconvoluting overlapping emission lines continue to evolve. We show here that quantitative fits of XFM data obtained after removing a baseline, calculated by connecting peak edges, can be unexpectedly dependent upon acquisition dwell-time and spectral aggregation leading to differences in apparent elemental content. Using mouse preimplantation embryos and ovarian follicles as model samples, we demonstrate how these variables influence quantitative comparisons between samples. We find that subtracting an empirically measured blank spectrum instead of a baseline provides quantitative XFM elemental mapping results that are independent of dwell time and spectral aggregation dependencies.

Supplemental Figures and Tables

| **Means** | *Si* | *P* | *S* | *Cl* | *Ar* | *K* | *Ca* | *Mn* | *Fe* | *Co* | *Cu* | *Zn* |
| --- | --- | --- | --- | --- | --- | --- | --- | --- | --- | --- | --- | --- |
| *Baseline (Aggregate)* | -17.2 | 4.73 | 5.17 | 36.6 | -0.137 | 9.89 | 0.274 | 3.63  $\times$ 10^-3^ | 1.25  $\times$ 10^-2^ | 1.64  $\times$ 10^-4^ | 3.79  $\times$ 10^-3^ | 5.98  $\times$ 10^-2^ |
| *Baseline (Per-Pixel)* | -13.6 | 7.59 | 7.09 | 41.4 | -0.131 | 10.8 | 0.313 | 7.37  $\times$ 10^-3^ | 1.49  $\times$ 10^-2^ | 2.53  $\times$ 10^-3^ | 5.34  $\times$ 10^-3^ | 6.06  $\times$ 10^-2^ |
| *Blank (Aggregate)* | -3.01 | 2.48 | 2.99 | 19.6 | 0.0407 | 7.09 | 0.241 | -1.81  $\times$ 10^-4^ | 1.39  $\times$ 10^-2^ | 5.04  $\times$ 10^-4^ | 4.36  $\times$ 10^-3^ | 6.64  $\times$ 10^-2^ |
| *Blank (Per-Pixel)* | -3.01 | 2.48 | 2.99 | 19.6 | 0.0407 | 7.09 | 0.241 | -1.81  $\times$ 10^-4^ | 1.39  $\times$ 10^-2^ | 5.04  $\times$ 10^-4^ | 4.36  $\times$ 10^-3^ | 6.64  $\times$ 10^-2^ |
| **Standard Deviations** | *Si* | *P* | *S* | *Cl* | *Ar* | *K* | *Ca* | *Mn* | *Fe* | *Co* | *Cu* | *Zn* |
| *Baseline (Aggregate)* | 5.05 | 0.988 | 0.822 | 7.10 | 3.28  $\times$ 10^-2^ | 2.34 | 8.98  $\times$ 10^-2^ | 1.10  $\times$ 10^-3^ | 4.37  $\times$ 10^-3^ | 1.33  $\times$ 10^-4^ | 1.07  $\times$ 10^-3^ | 1.53  $\times$ 10^-2^ |
| *Baseline (Per-Pixel)* | 5.85 | 1.27 | 1.07 | 7.56 | 3.92  $\times$ 10^-2^ | 2.70 | 9.35  $\times$ 10^-2^ | 1.96  $\times$ 10^-3^ | 5.08  $\times$ 10^-3^ | 6.31  $\times$ 10^-4^ | 1.35  $\times$ 10^-3^ | 1.65  $\times$ 10^-2^ |
| *Blank (Aggregate)* | 1.18 | 0.403 | 0.418 | 3.39 | 2.28  $\times$ 10^-2^ | 1.64 | 7.05  $\times$ 10^-2^ | 3.88  $\times$ 10^-4^ | 4.62  $\times$ 10^-3^ | 9.81  $\times$ 10^-5^ | 1.14  $\times$ 10^-3^ | 1.68  $\times$ 10^-2^ |
| *Blank (Per-Pixel)* | 1.18 | 0.403 | 0.418 | 3.39 | 2.28  $\times$ 10^-2^ | 1.64 | 7.05  $\times$ 10^-2^ | 3.88  $\times$ 10^-4^ | 4.62  $\times$ 10^-3^ | 9.81  $\times$ 10^-5^ | 1.14  $\times$ 10^-3^ | 1.68  $\times$ 10^-2^ |

**Table S1:** The mean quantification and population distribution in units of µg/cm^2^ from the ROI analysis of the 12 blastocysts shown in Fig. 1 resulting from baseline subtraction and blank spectrum correction of per-pixel and aggregate spectra. Comparison of blank spectrum correction for aggregate and per-pixel spectra shows the answers are identical and therefore independent of any degree of signal aggregation.

|  | **Blank (Aggregate vs. Per-Pixel)** | | | |
| --- | --- | --- | --- | --- |
|  | **Δ Conc.** | **Δ Pop. Dist.** | **Paired T-Test** | **Unpaired T-Test** |
| *Si* | -3.9 $\times$10^-11^ % | 3.21 $\times$10^-11^ % | #DIV/0 | 1 |
| *P* | -2.56 $\times$10^-10^ % | -5.23 $\times$10^-10^ % | #DIV/0 | 1 |
| *S* | -5.56 $\times$10^-11^ % | -5.65 $\times$10^-10^ % | 0.999146 | 1 |
| *Cl* | 1.82 $\times$10^-10^ % | 4.62 $\times$10^-10^ % | #DIV/0 | 1 |
| *Ar* | 2.4 $\times$10^-11^ % | -7.58 $\times$10^-11^ % | 0.99992 | 1 |
| *K* | -1.14 $\times$10^-10^ % | 6.15 $\times$10^-10^ % | #DIV/0 | 1 |
| *Ca* | -1.78 $\times$10^-10^ % | -1.58 $\times$10^-10^ % | #DIV/0 | 1 |
| *Mn* | -1.05 $\times$10^-9^ % | -5.43 $\times$10^-11^ % | 0.9999 | 1 |
| *Fe* | -1.82 $\times$10^-10^ % | -1.10 $\times$10^-10^ % | #DIV/0 | 1 |
| *Co* | 2.14 $\times$10^-10^ % | -1.37 $\times$10^-10^ % | #DIV/0 | 1 |
| *Cu* | 7.85 $\times$10^-11^ % | 7.33 $\times$10^-11^ % | 0.999551 | 1 |
| *Zn* | 9.45 $\times$10^-11^ % | 1.53 $\times$10^-10^ % | 0.999273 | 1 |

**Table S2:** Relative percentage differences in quantification and population distribution from ROI analysis of the 12 blastocysts shown in Fig. 1, and tabulated in Table S1, resulting from blank spectrum correction of per-pixel and aggregate spectra. Statistical differences were determined using a paired-sample t-test, with the results of unpaired-sample t-test shown as a courtesy.

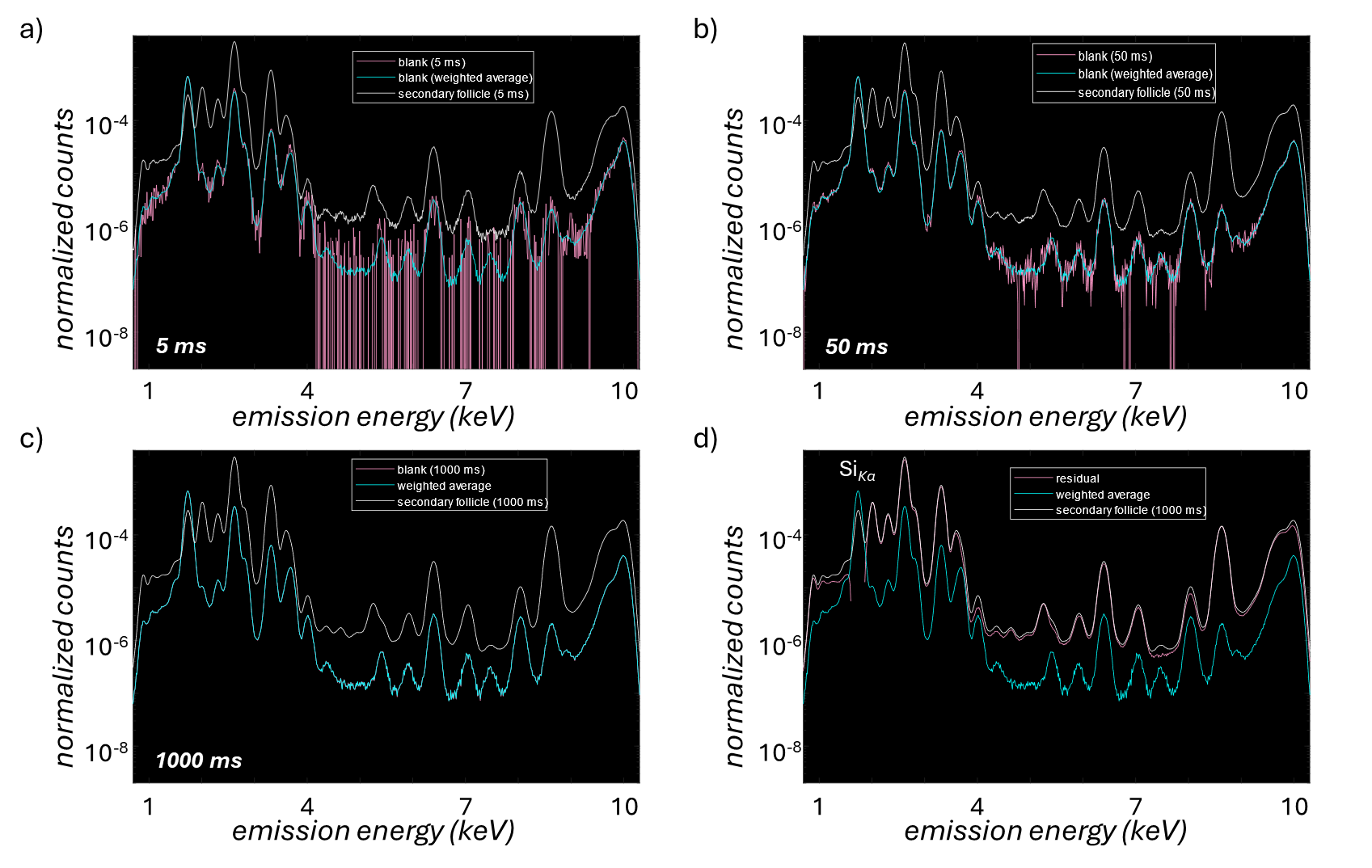

**Figure S1**: Blanks used in blank spectrum correction of the secondary follicle presented in Figure 2 of the main text. X-ray fluorescence microscopy mapping of a subregion of a murine secondary ovarian follicle was performed at dwell times of 5 ms, 10 ms, 50 ms, and 1000 ms with subsequent background correction performed in M-BLANK using the blank spectrum correction approach. (a-c) show the calculated blanks from the different dwell times of 5 ms, 50 ms, and 1000 ms are shown along with the blank obtained from the weighted average of all three. d) shows the residual signal post blank spectrum correction of the 1000 ms dwell time image corrected using the weighted average blank.

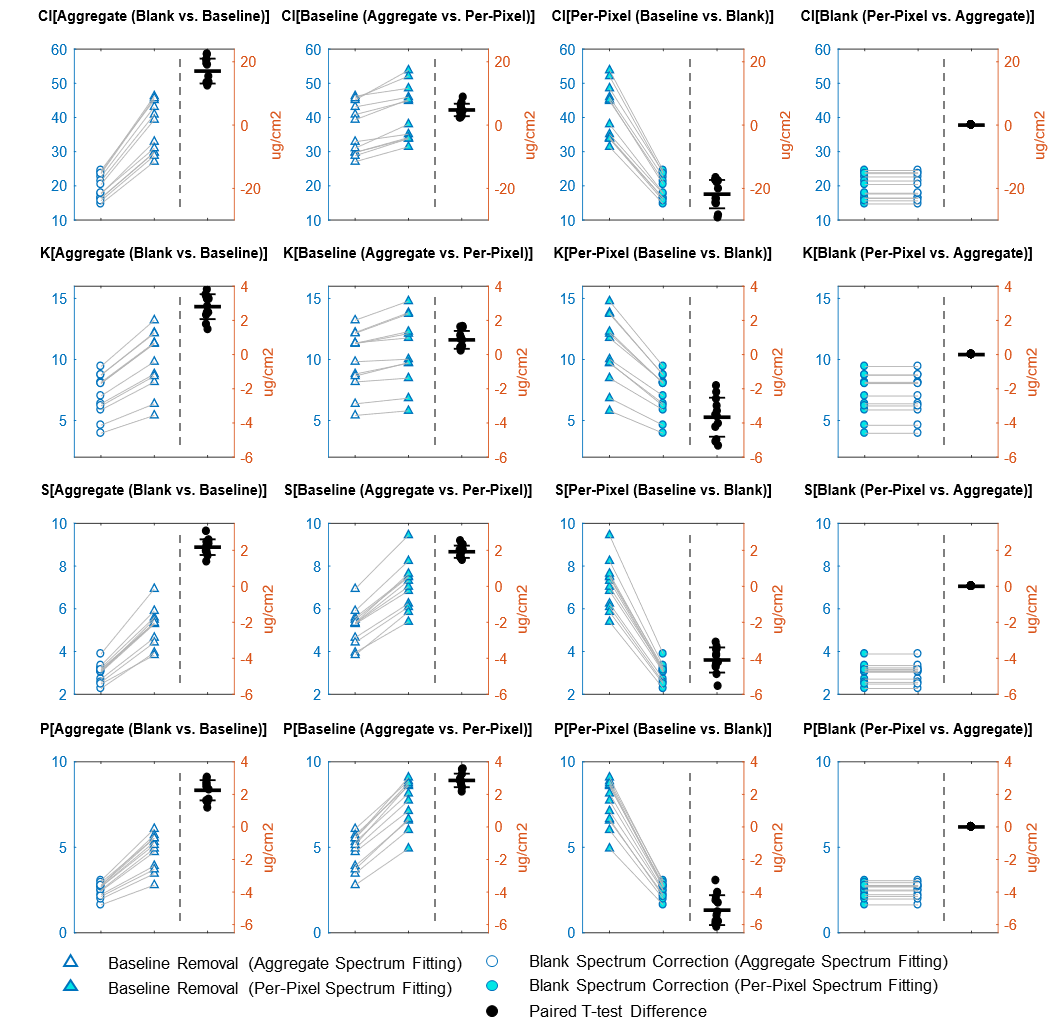

**Figure S2:** Pair-wise plots from paired-sample t-tests performed on the 12 blastocysts from Fig. 1 for chlorine, potassium, sulfur, and phosphorus. Statistical differences were determined using a paired-sample t-test: * p<0.05; ** p<0.01; *** p<0.001; **** p<0.0001.

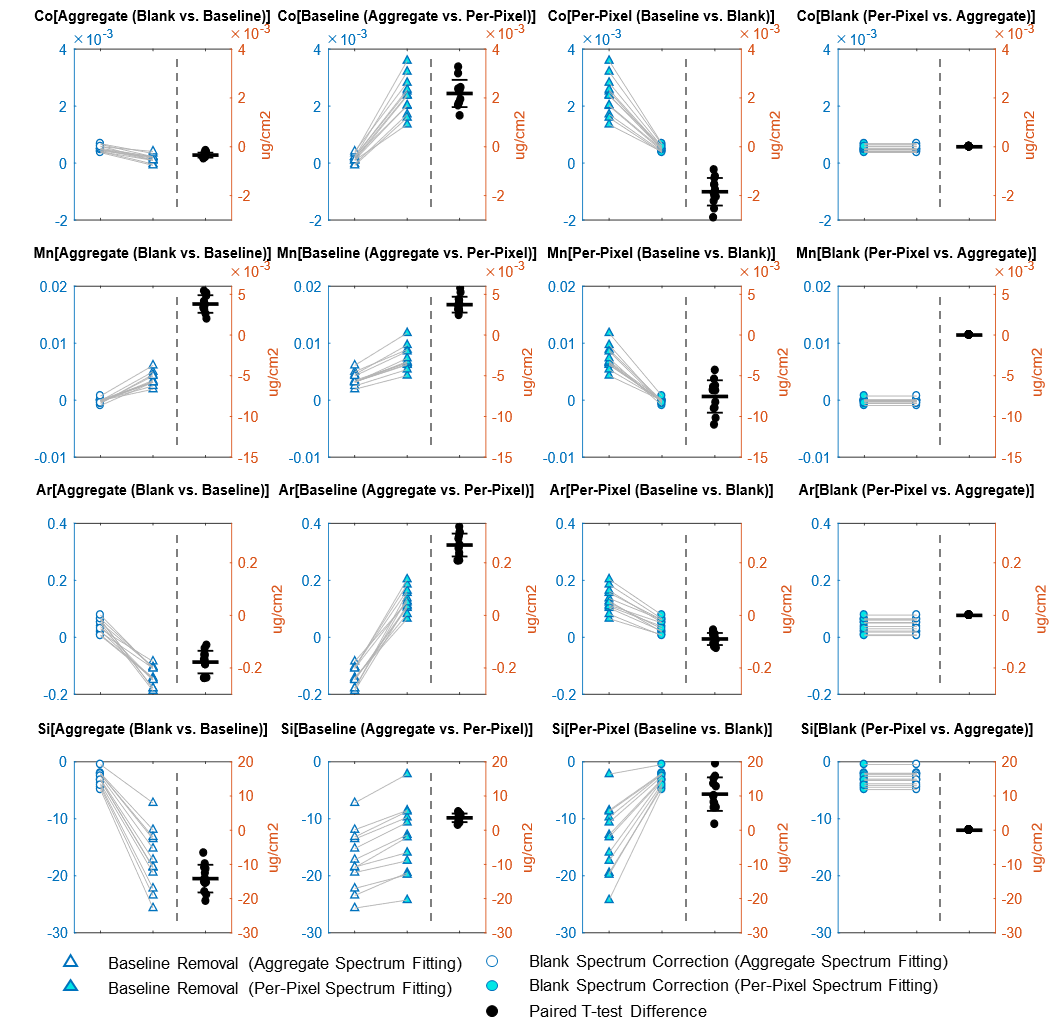

**Figure S3:** Pair-wise plots from paired-sample t-tests performed on the 12 blastocysts from Fig. 1 for calcium, zinc, iron, and copper. Statistical differences were determined using a paired-sample t-test: * p<0.05; ** p<0.01; *** p<0.001; **** p<0.0001.

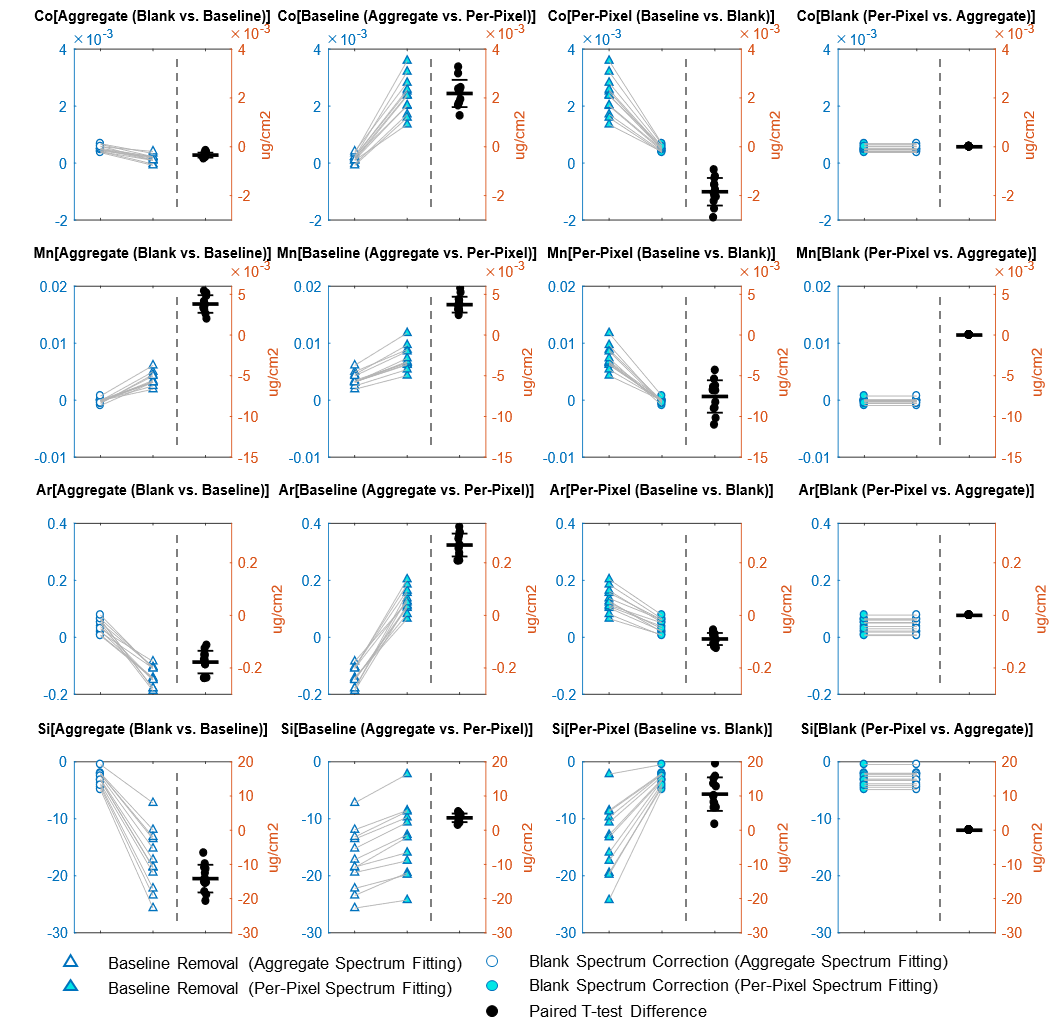

**Figure S4:** Pair-wise plots from paired-sample t-tests performed on the 12 blastocysts from Fig. 1 for cobalt, manganese, ambient argon, and substrate silicon. Statistical differences were determined using a paired-sample t-test: * p<0.05; ** p<0.01; *** p<0.001; **** p<0.0001.

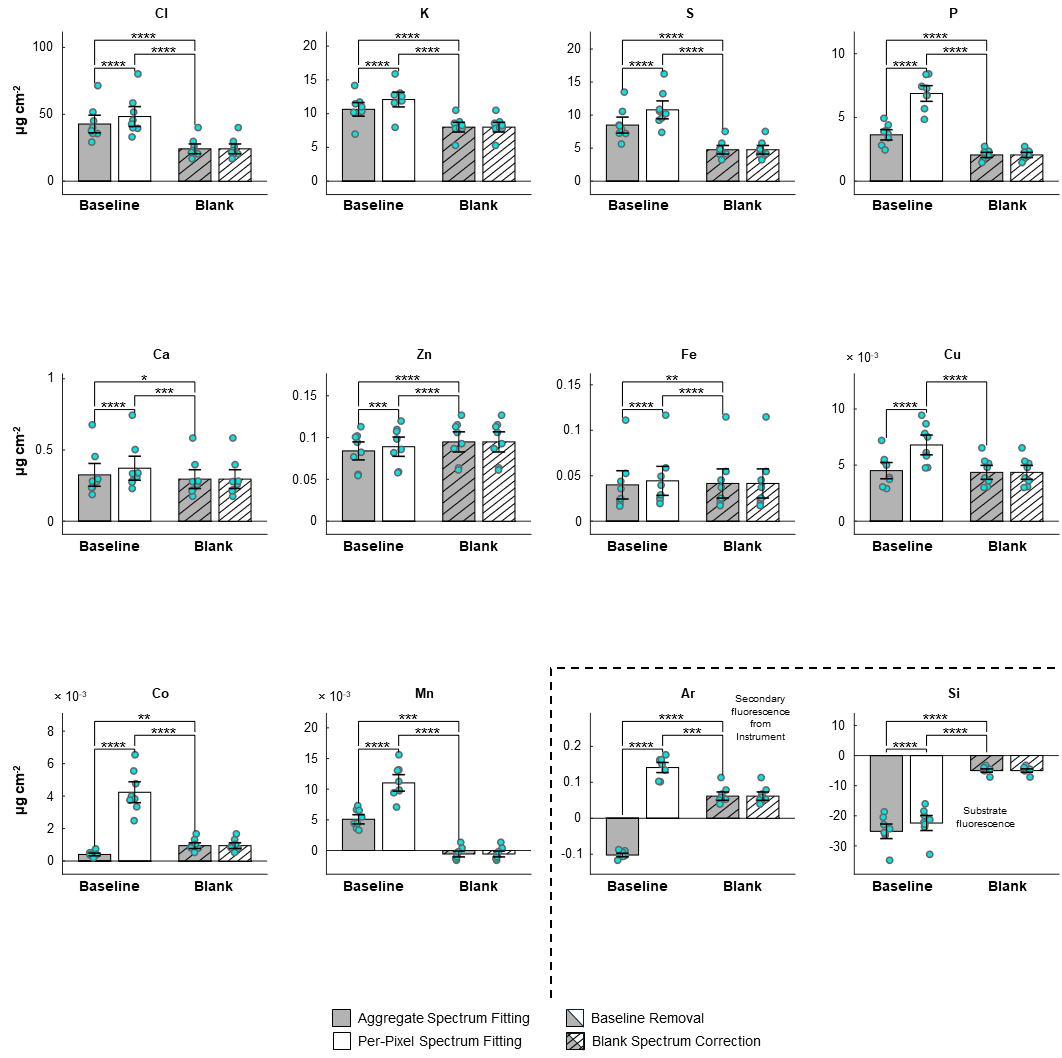

**Figure S5:** Differences in quantitative analysis of early-stage blastocysts between baseline and blank spectrum correction of per-pixel and aggregate spectra. When analyzing the concentration of relevant elements across a set of 8 early stage blastocytes, significant quantitative differences exist depending on whether baseline or blank background subtraction methods were used and whether analysis was performed on individual pixel spectra or the aggregate spectrum of image ROIs. Quantitatively, blank spectrum correction fitting of per-pixel and aggregate spectra differed negligibly with relative root-mean-squared-errors ca. 1×10^-14^. However, baseline subtraction of per-pixel and aggregate spectra never agreed, nor did either agree with quantitation from blank spectrum correction fitting. Bar plots correspond to the population mean. Errors bars are set at one standard deviation. Statistical differences were determined using a paired-sample t-test: * p<0.05; ** p<0.01; *** p<0.001; **** p<0.0001.

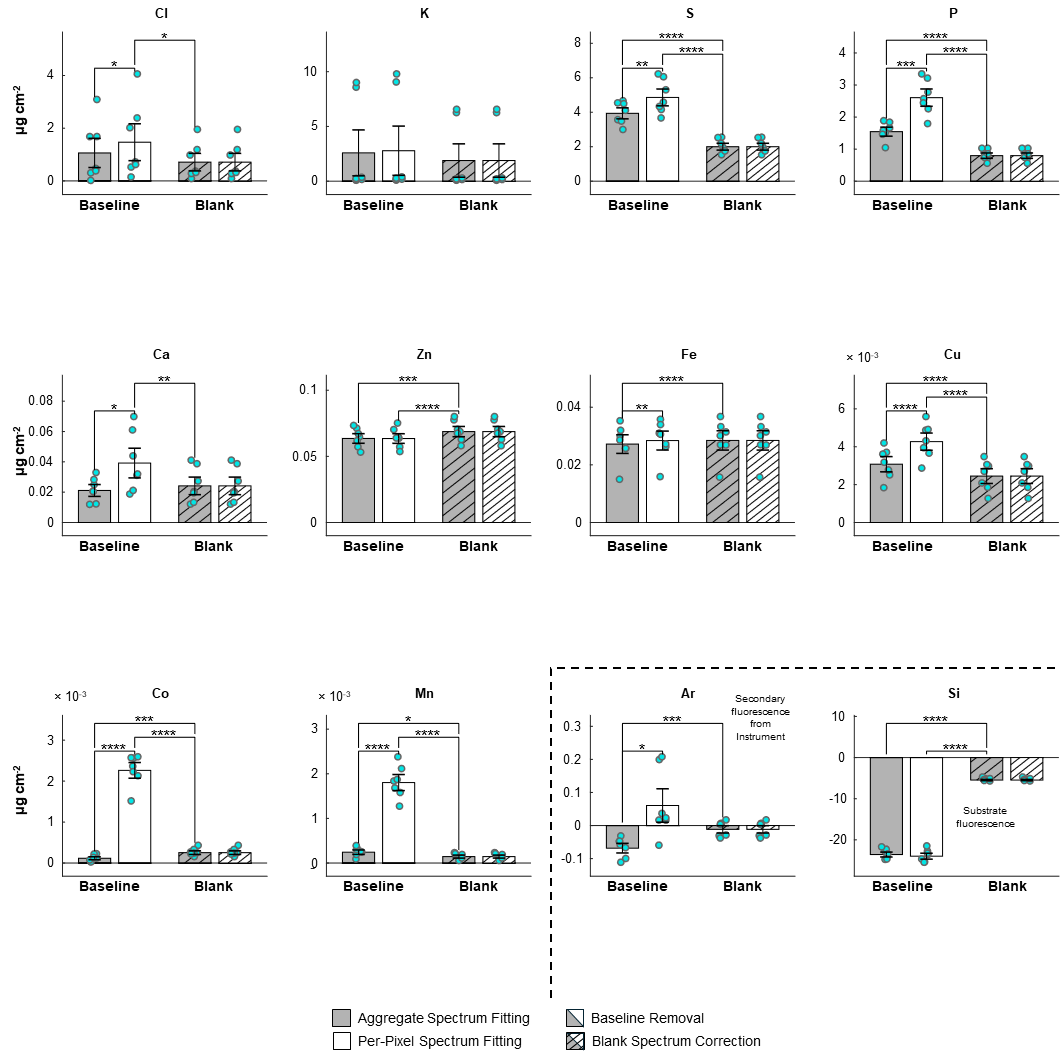

**Figure S6:** Comparison of 8-cell Embryo - Differences in quantitative analysis of 8-cell stage embryos between baseline and blank spectrum correction of per-pixel and aggregate spectra. When analyzing the concentration of relevant elements across a set of 7 8-cell stage embryos, significant quantitative differences exist depending on whether baseline or blank background subtraction methods were used and whether analysis was performed on individual pixel spectra or the aggregate spectrum of image ROIs. Quantitatively, blank spectrum correction fitting of per-pixel and aggregate spectra differed negligibly with relative root-mean-squared-errors ca. 1×10^-14^. However, baseline subtraction of per-pixel and aggregate spectra never agreed, nor did either agree with quantitation from blank spectrum correction fitting. Bar plots correspond to the population mean. Errors bars are set at one standard deviation. Statistical differences were determined using a paired-sample t-test: * p<0.05; ** p<0.01; *** p<0.001; **** p<0.0001.

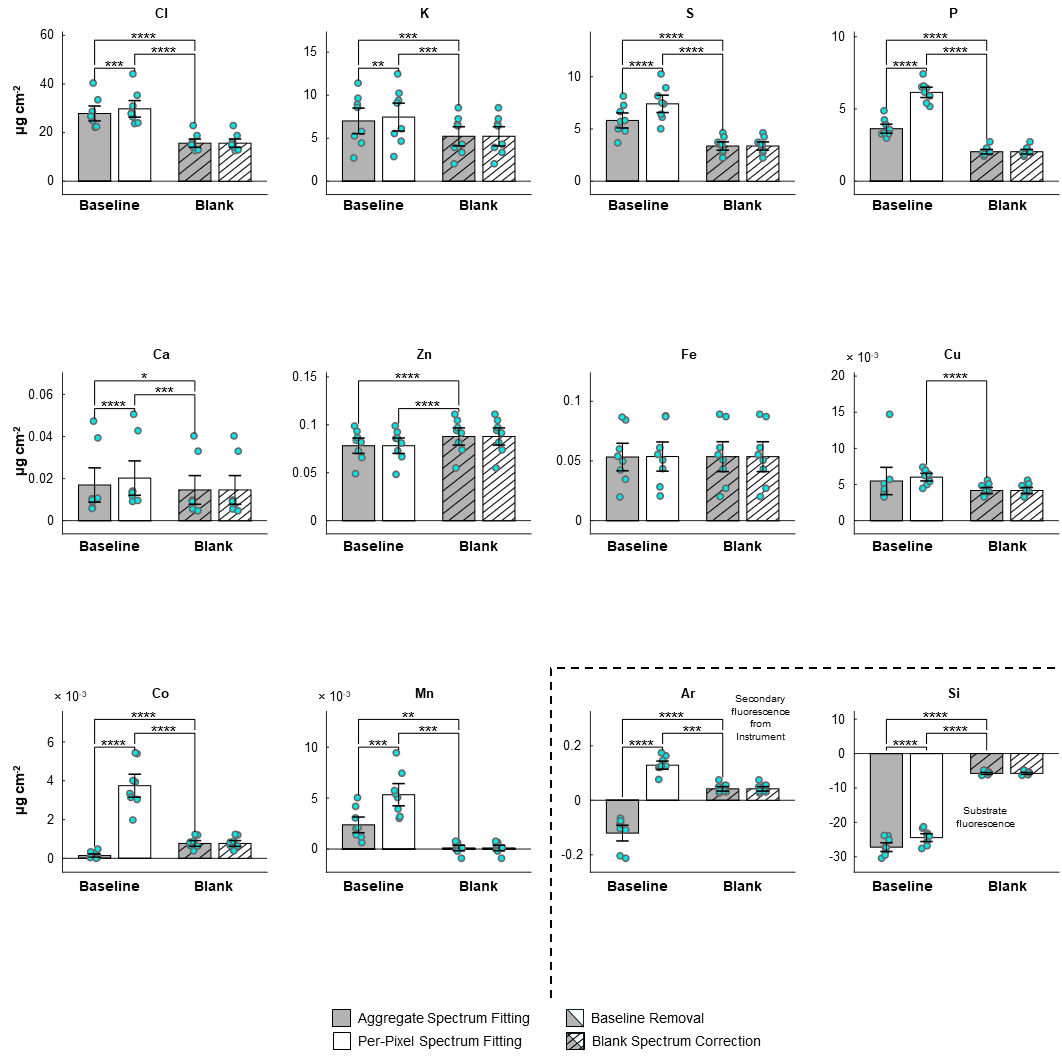

**Figure S7:** Comparison of Morula Differences in quantitative analysis of 8 morula stage embryos between baseline and blank spectrum correction of per-pixel and aggregate spectra. When analyzing the concentration of relevant elements across a set of 7 8-cell stage embryos, significant quantitative differences exist depending on whether baseline or blank background subtraction methods were used and whether analysis was performed on individual pixel spectra or the aggregate spectrum of image ROIs. Quantitatively, blank spectrum correction fitting of per-pixel and aggregate spectra differed negligibly with relative root-mean-squared-errors ca. 1×10^-14^. However, baseline subtraction of per-pixel and aggregate spectra never agreed, nor did either agree with quantitation from blank spectrum correction fitting. Bar plots correspond to the population mean. Errors bars are set at one standard deviation. Statistical differences were determined using a paired-sample t-test: * p<0.05; ** p<0.01; *** p<0.001; **** p<0.0001.

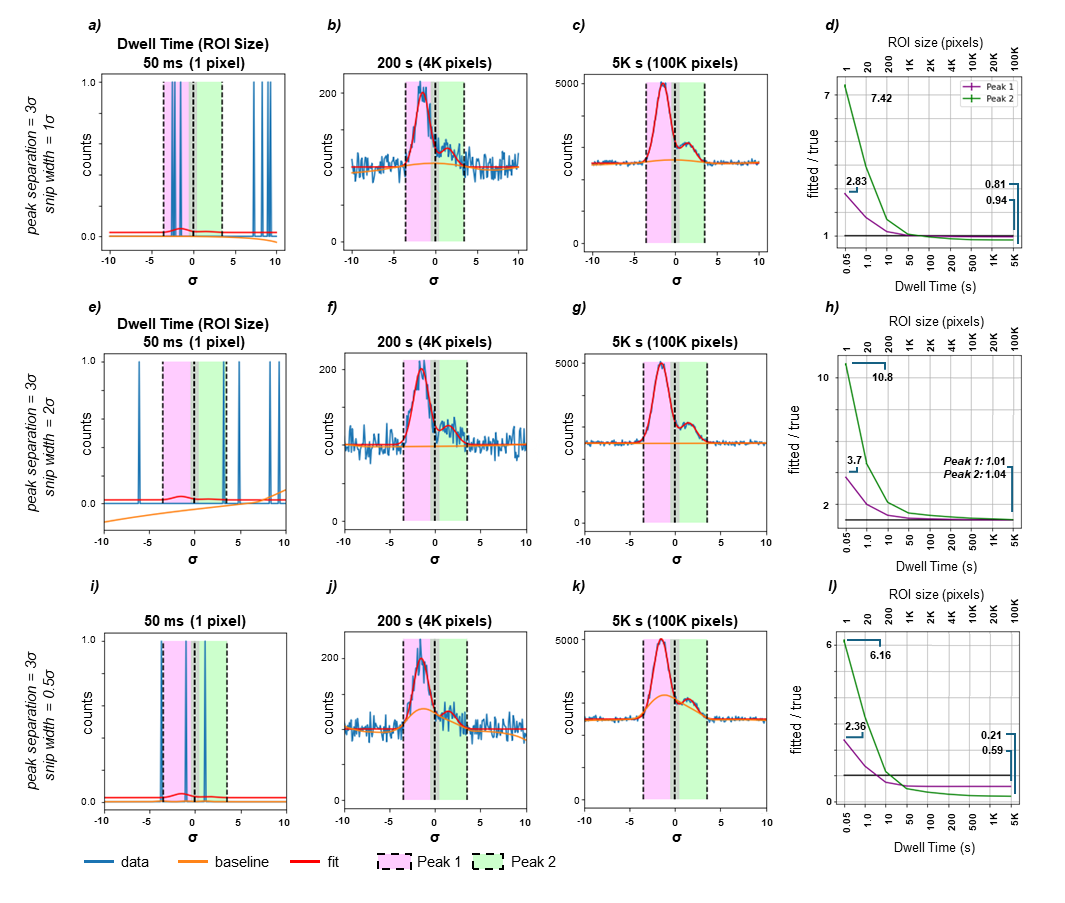

**Figure S8**: Baseline distortion is also systematically dependent on the width of the SNIP function. We used the same model from Figure 6 consisting of two Gaussian functions riding on a constant linear background to explore the apparent peak area dependencies on the snip-width as a function of statistical noise, calculated from a Poisson distribution. The model was defined to have a signal-to-noise ratio of 1 for 1 second. We started with a single Gaussian with peak height of 1 and a background of 1. To study the effect of peak overlap, a second, smaller peak (peak height = 0.25) was added. Both Gaussians had a full-width half-maximum of 1 and were separated by 3 (a-e) and 6 (f-j) standard deviation units. Simulations were done for dwell times ranging from 50 milliseconds to 5,000 seconds (effective image ROI sizes of 1 pixel to 100,000 pixels), with representative simulations for 50 milliseconds (a, f, and i), 200 seconds (b, g, and j), and 5,000 seconds (c, h, and k) shown. For each of the resulting data sets, baseline subtraction was modeled as a SNIP baseline calculated using the pybaselines implementation of the SNIP algorithm (<https://pypi.org/project/pybaselines/>) with 5 point moving-average smooth (keyword smooth_half_window = 2) and a snip-width (keyword half_max_width) set at multiples of the full-width-at-half-max of 1σ (a-d), 2σ (e-h), and 0.5σ (i-l). Blank spectrum correction was not modeled because there is no snip-width parameter. In order to estimate the reliability of the simulations, each condition was repeated 10,000 times and the resulting ratios were averaged to give the mean and standard deviation.

Python code for simulations:

#!/usr/bin/env python

\# coding: utf-8

\# Original description of SNIP is: Ryan et al, NIM, 1988 : https://www.sciencedirect.com/science/article/pii/0168583X88900638

\#

\# Impplementation used here is from PyBaselines: see `https://pypi.org/project/pybaselines/`

\# To use pybaselines, need to install it

\# `conda install -c conda-forge pybaselines`

\# In[19]:

import numpy as np

import matplotlib.pyplot as plt

import pybaselines

def gauss(fwhm,center,x):

sigma = fwhm/(2*np.sqrt(np.log(2)))

return np.exp( -((x-center)/sigma)**2) #For peakheight of 1

def run_sim(spacing,amplitude=1,fwhm=2,use_blank_correction = False):

#The parameters is this first block will change the type of calculation that is done

\# fwhm = 2 #FWHM for the Gaussians

N_gauss = 2 #This can be set > 2 providing g_cent and g_amp are increased. However, titles on plots won't work

\# spacing=g_cent[1]-g_cent[0]

\# spacing=3

g_cent = [-spacing/2,spacing/2]

#g_cent = [-4,4]

g_amp = np.array((1,0.25))*amplitude

xmax=max(10,spacing)

xmin=-1*xmax

\# xmin = -10. #Minimum x value - this should be adjusted to match the gaussian peaks

\# xmax = 10. #Maximum x value

delta = 0.1 #Point spacing in x

dwellTimes = [0.05, 1, 10, 50, 100, 200, 500, 1000, 5000]#[3,10,100,200,500,1000,2000,5000,10000] #Data (below) have a maximum at the peak of 1 counts/second

\# dwellTimes = [10000,5000,2000,1000,500,200,100,10,3] #Data (below) have a maximum at the peak of 1 counts/second

S_N = 1. #This is the ratio of the height of the standard gaussian (before g_amp scaling) / size of linear background

#This means that S_N = 1 corresponds to 0.5 signal counts/second and 0.5 background counts. S_N=9 is 0.9 signal/sec and

#0.1 background/sec.

n_statistics = 10000 #10000 #Number of times to repeat the calculation to get better statistics.

#----------------------------------------

#Define ROIs for integration and signal

A_gauss = S_N/(1+S_N)

A_back = 1./(1+S_N)

npt_w = int(fwhm/delta) #Parameter needed for SNIP - data points correspond to fwhm

npt_s = 2 #Half-window for running smooth in SNIP

x = np.arange(xmin,xmax + delta/2.,delta) #Add the delta/2 to avoid round-off errors

ROI = np.zeros((len(x),N_gauss), dtype = bool) #Holder for filters to calculate ROIs for each peak.

signal = np.zeros(len(x))

#Determine the limits

ROI_lims = np.zeros((N_gauss,2))

for ng in range(N_gauss):

ROI_lims[ng,0] = g_cent[ng] - fwhm

ROI_lims[ng,1] = g_cent[ng] + fwhm

#Trim if necessary to avoid overlap

ROI_lims_orig=ROI_lims+0

for ng in range(N_gauss-1):

upperN = ROI_lims[ng,1]

lowerN1 = ROI_lims[ng+1,0]

if upperN > lowerN1: #Go through the ROIs. If the upper value for N is larger than the lower value for N+1, average them

newLim = (upperN+lowerN1)/2.

ROI_lims[ng,1] = newLim

ROI_lims[ng+1,0] = newLim

for ng in range(N_gauss):

ROI[:,ng] = np.logical_and(x >= ROI_lims[ng,0], x <= ROI_lims[ng,1]) #Filter for ROI - define as 2 x fwhm

signal += A_gauss * g_amp[ng] * gauss(fwhm,g_cent[ng],x)

background = A_back

y = signal + background

y_n = np.zeros(len(y))

#----------------------------------------------

#Set up arrays to hold the data when it's calculated, calculate true areas, set up plots

baselines = np.zeros((len(x))) #To hold the baselines

correctedSignal = np.zeros((len(x))) #Signal after baseline subtraction

ratios = np.zeros((len(dwellTimes),n_statistics,N_gauss)) #Third dimension is for each peak's ROI

trueArea = np.zeros(N_gauss) #Determine the areas that each peak "should" have, for an ROI that goes from -fwhm to + fwhm around center

for ng in range(N_gauss):

trueArea[ng] = np.sum(signal[ROI[:,ng]])

nrows = 3 # Ros in the plot for different times

ncols = int(np.ceil(len(dwellTimes)/nrows))

fig,axn = plt.subplots(nrows = nrows, ncols = ncols, figsize = (12,4*nrows)) #Create a figure to show sample data

ax = np.ravel(axn)

#--------------------------------------------

#Calculate the data for each dwell time. Plot the first set of data as representative

colors=['purple','green']

fcs=0.2

fcs=[(1,0,1,fcs),(0,1,0,fcs)]

for n_d, dwell in enumerate(dwellTimes):

for count in range(n_statistics): #Execute n_statistics times to get statistics for ratio

for n, yv in enumerate(y):

y_n[n] = np.random.poisson(yv*dwell)

if not use_blank_correction:

baselines[:] = pybaselines.smooth.snip(y_n, max_half_window=npt_w, decreasing=True, smooth_half_window= npt_s)[0]

correctedSignal[:] = y_n - baselines[:]

else:

nsamp4blank=1e6*0.05

baselines[:] = np.random.normal(background *nsamp4blank, np.sqrt(background *nsamp4blank))/nsamp4blank * dwell

correctedSignal[:] = y_n - baselines[:] # subtract the simulated blank

for nROI in range(N_gauss):

ratios[n_d,count,nROI] = np.sum(correctedSignal[ROI[:,nROI]])/trueArea[nROI]/dwell

if count == 0: #Only plot the first data set

ax[n_d].plot(x,y_n, label = 'signal')

ax[n_d].plot(x,y*dwell, color = 'red')

ax[n_d].plot(x,baselines[:])

ax[n_d].set_title('Dwell = {:}'.format(dwell))

for nvert in range(N_gauss):

mx=max((dwell,max(y_n)))

ax[n_d].plot(ROI_lims[nvert,0]*np.ones(2),[0,mx],color = 'black', linestyle = 'dashed')

ax[n_d].plot(ROI_lims[nvert,1]*np.ones(2),[0,mx],color = 'black', linestyle = 'dashed')

pp1 = plt.Rectangle((ROI_lims_orig[nvert,0],0),fwhm*2,mx,fc=fcs[nvert])

ax[n_d].add_patch(pp1)

plt.suptitle('Spacing = {:.1f}, fwhm = {:.1f}, Amp ratio = {:.1f}, S/B = {:.1f}, max_window = {:.1f}, smooth_window = {:.1f}'

.format(spacing, fwhm,g_amp[1]/g_amp[0],amplitude,(2*npt_w+1)*delta, (2*npt_s+1)*delta))

#plt.suptitle('Spacing = {:.1f}, fwhm = {:.1f}, Amp ratio = {:.1f}, max_window = {:.1f}, smooth_window = {:.1f}'

\#	.format(g_cent[1]-g_cent[0], fwhm,g_amp[1]/g_amp[0],(2*npt_w+1)*delta, (2*npt_s+1)*delta))

plt.show()

#-----------------------------------------

#Plot statistics

aveRatio = np.average(ratios, axis =1)

aveStd = np.std(ratios, axis =1)

tickText = []

for d in dwellTimes:

tickText.append('{:.02f}'.format(d))

fig,ax = plt.subplots(figsize = (4,4))

for ng in range(N_gauss):

ax.errorbar(np.arange(len(dwellTimes)),y = aveRatio[:,ng], yerr = aveStd[:,ng]/np.sqrt(n_statistics),

label = 'Peak {}'.format(ng+1), color=colors[ng])

ax.plot(np.arange(len(dwellTimes)),np.ones(len(dwellTimes)), color = 'black')

ax.set_ylabel('Baseline subtracted counts/true counts')

ax.set_xticks(np.arange(len(dwellTimes)))

ax.set_xticklabels(tickText, rotation = 'vertical')

ax.set_xlabel('Dwell time')

\# ax.set_ylim([0.5, 10])

ax.legend()

ax.grid()

ax.set_title('Area ratios for different dwell times')

plt.show()

\# In[ ]:

run_sim(3)

run_sim(6)

\# run_sim(3,.2)

\# run_sim(6,.2)

run_sim(3,use_blank_correction = True)

run_sim(6,use_blank_correction = True)
