## Supplementary figures and images for "Blank Spectrum Correction as a Robust Solution to Artifacts in Quantitative X-ray Fluorescence Mapping"

### Table of Content Graphic

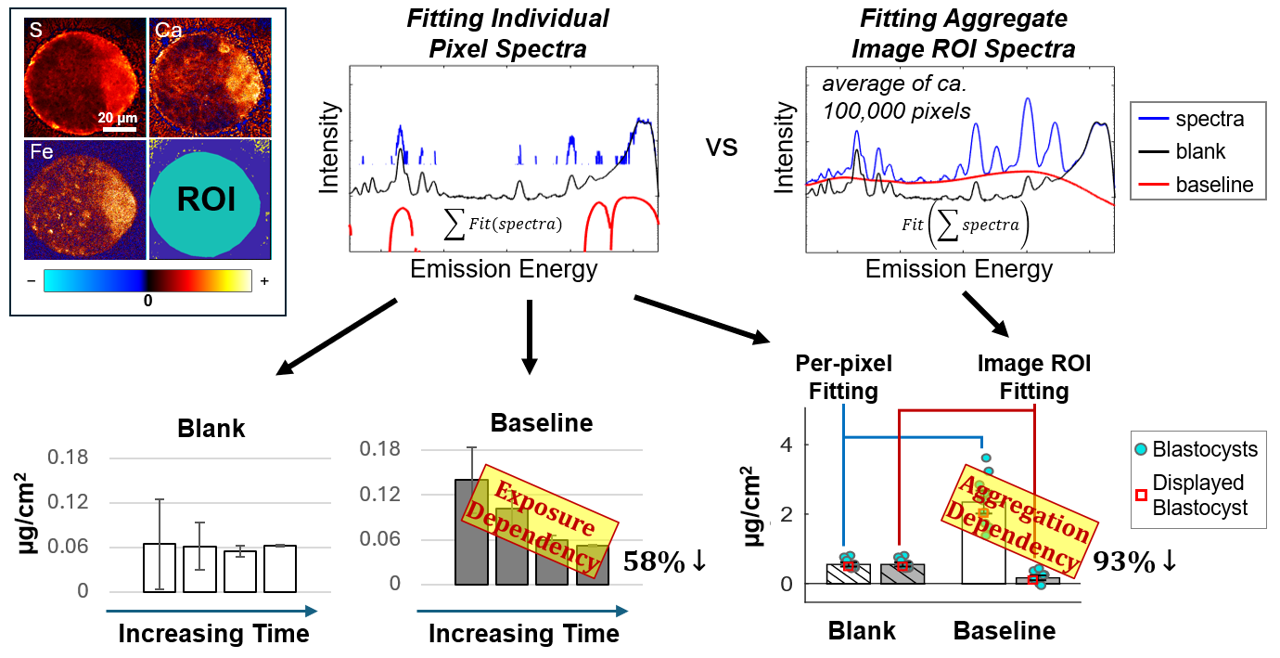
